# Identification of genetic variants that impact gene co-expression relationships using large-scale single-cell data

**DOI:** 10.1101/2022.04.20.488925

**Authors:** Shuang Li, Katharina T. Schmid, Dylan de Vries, Maryna Korshevniuk, Roy Oelen, Irene van Blokland, BIOS Consortium, sc-eQTLgen Consortium, Hilde E. Groot, Morris Swertz, Pim van der Harst, Harm-Jan Westra, Monique van der Wijst, Matthias Heinig, Lude Franke

## Abstract

**Background:** Expression quantitative trait loci (eQTL) studies have shown how genetic variants affect downstream gene expression. To identify the upstream regulatory processes, single-cell data can be used. Single-cell data also offers the unique opportunity to reconstruct personalized co-expression networks—by exploiting the large number of cells per individual, we can identify SNPs that alter co-expression patterns (co-expression QTLs, co-eQTLs) using a limited number of individuals.

**Results:** To tackle the large multiple testing burden associated with a genome-wide analysis (i.e. the need to assess all combinations of SNPs and gene pairs), we conducted a co-eQTL meta-analysis across four scRNA-seq peripheral blood mononuclear cell datasets from three studies (reflecting 173 unique participants and 1 million cells) using a novel filtering strategy followed by a permutation-based approach. Before analysis, we evaluated the co-expression patterns to be used for co-eQTL identification using different external resources. The subsequent analysis identified a robust set of cell-type-specific co-eQTLs for 72 independent SNPs that affect 946 gene pairs, which we then replicated in a large bulk cohort. These co-eQTLs provide novel insights into how disease-associated variants alter regulatory networks. For instance, one co-eQTL SNP, rs1131017, that is associated with several autoimmune diseases affects the co-expression of *RPS26* with other ribosomal genes. Interestingly, specifically in T cells, the SNP additionally affects co-expression of *RPS26* and a group of genes associated with T cell-activation and autoimmune disease. Among these genes, we identified enrichment for targets of five T-cell-activation-related transcriptional factors whose binding sites harbor rs1131017. This reveals a previously overlooked process and pinpoints potential regulators that could explain the association of rs1131017 with autoimmune diseases.

**Conclusion:** Our co-eQTL results highlight the importance of studying gene regulation at the context-specific level to understand the biological implications of genetic variation. With the expected growth of sc-eQTL datasets, our strategy—combined with our technical guidelines—will soon identify many more co-eQTLs, further helping to elucidate unknown disease mechanisms.

## Background

In recent years, genome-wide association studies (GWAS) have revealed a large number of associations between genetic variation and disease (1). Many of these variants also change downstream gene expression, as identified using expression quantitative trait locus (eQTL) analysis (2). However, even with many such connections now identified, the upstream biological processes that regulate these eQTLs often remain hidden. Such knowledge is important for better understanding the underlying processes that lead to specific disease, which would aide in drug development (3).

One way to study the biological processes in which eQTL genes are involved is to construct gene co-expression networks. In these networks, genes (nodes) involved in shared biological processes are expected to be connected through co-expression (edges) (4). Traditionally, these networks have been reconstructed with bulk RNA sequencing (RNA-seq) data, using a variety of computational tools (5–7). However, whether certain biological processes are active can depend on various factors, such as cell type, environmental exposures and even single nucleotide polymorphisms (SNPs) (2,8,9). With single-cell technologies, many of these highly specific contexts can now be captured at high resolution. Single-cell RNA-seq (scRNA-seq) not only allows for cell-type-specific analyses, it does so without the technical biases introduced by the cell-sorting required to perform similar analyses with bulk RNA-seq.

In addition to capturing the cell-type-specific contexts, scRNA-seq can also be used to construct personalized co-expression networks using the repeated measurements (i.e. multiple single-cell gene expression profiles) for each individual. This enables quantification of the covariance between genes, and thus their co-expression strengths, within an individual (10). These personalized co-expression networks can then be used to study the effects of genetic variation on network properties. Some of these network changes can be linked to individual SNP genotypes, called co-expression quantitative trait loci (co-eQTLs).

While we have previously shown that co-eQTLs can be both cell-type-specific and stimulation-specific, several challenges to systematic identification remain (10, 11). Firstly, it is unclear how to best construct gene regulatory networks (GRNs) with scRNA-seq data. Co-expression patterns identified from bulk RNA sequencing data have been shown to be informative for physical and functional gene–gene interactions (5–7), but it is unclear whether the co-expression patterns identified with scRNA-seq data also reflect gene–gene functional interactions given technical challenges of scRNA-seq data such as sparseness and low signal-to-noise ratios (12, 13). These issues are caused by a combination of low mRNA counts in cells, imperfect capture efficiencies and the inherent stochasticity of mRNA expression (14). Many methods have been proposed to account for this issue. A recent benchmark paper (15) suggested ‘rho proportionality’ (16) as an association measure because of its consistent performance. Also complementary strategies could be beneficial, such as combining association measures with *MetaCell,* a recently proposed method that groups homogeneous cells to reduce sparsity, but to our knowledge it has not yet been evaluated in benchmark studies (17). Moreover, a recent benchmark paper concluded that different GRN construction methods show moderate performance that is often dataset-specific (18), indicating that many challenges remain in GRN reconstruction. Therefore, validation of the robustness and functional relevance of the network is warranted.

Secondly, there is no consensus method for co-eQTL mapping and personalized GRN construction. In bulk data with only one measurement per individual, it is not possible to identify co-eQTLs directly. To carry out a similar type of analysis in bulk data, we previously used a linear regression model with an interaction term to identify interaction QTLs in bulk data from whole blood (8). This approach can reveal co-eQTLs using the expression levels of individual genes as interaction terms. However, as bulk data nearly always comprises a mixture of cell types, it is not straightforward to unequivocally conclude that eQTLs showing an interaction effect reflect co-eQTLs (genetic variants that affect the co-expression between pairs of genes). A further compounding problem is that very large numbers of samples are required to identify co-eQTLs, and effects that manifest in specific (rare) cell types can easily be missed because they are masked by more common cell types. In theory, single-cell data allows direct estimation of cell-type-specific and individual-specific co-expression strength and should reduce the sample size requirement compared to bulk datasets. However, in practical terms, there are currently no datasets large enough to provide the statistical power to do genome-wide co-eQTL mapping, as this involves a large multiple testing burden due to billions of tests for every SNP and every possible gene pair combination. As such, there is a clear need for a robust co-eQTL strategy that can overcome the severe multiple testing issues and deal with the aforementioned issues with regards to the construction of reliable personalized co-expression networks.

In this work, we studied the genetic regulation of gene co-expression by conducting the largest-to-date co-eQTL meta-analysis in 173 peripheral blood mononuclear cell (PBMC) scRNA-seq samples. Before conducting this co-eQTL analysis, we determined the best strategy to identify cell-type-specific co-expression relationships in scRNA-seq data by benchmarking various methods and studying them in several independent datasets, including bulk RNA-seq and a CRISPR-coupled scRNA-seq screen knockout dataset. We then studied the effects of cell-type and inter-individual differences in gene co-expression networks by reconstructing personalized and cell-type-specific networks. We subsequently developed a robust co-eQTL mapping strategy with a novel filtering approach and an adapted permutation-based multiple testing procedure to deal with the correlation structure in the co-expression networks. By applying this strategy, we could perform a co-eQTL meta-analysis using data from three different scRNA-seq studies. We provided a comprehensive analysis of the different factors that affect the quality and quantity of co-eQTLs, including the number of cells, gene expression levels and filtering strategy. We then studied which biological processes and genes are regulated by the identified co-eQTLs by performing different enrichment analyses and exploring common biological functions, transcription factor (TF) binding and disease associations to try and pinpoint potential direct regulators of the co-eQTL genes. In sum, our results suggest that the combination of a robust method and a large sample size is crucial for identification of genetic variants that affect co-expression networks.

## Results

### Overview of the study

To uncover the contexts and biological processes that affect gene expression regulation, this study took advantage of both the resolution of single-cell data and the directionality captured by co-eQTLs. First, we constructed cell-type-specific co-expression networks from three recently generated PBMC scRNA-seq studies totaling 187 individuals and approximately one million cells. In two of the studies, donors were measured using two different versions of 10X Genomics chemistry (version 2 or version 3). To avoid batch effects due to these technical differences, we split both studies into two datasets, depending on the chemistry, leading to five datasets in total: 1) two datasets from the Oelen study (11) that collected unstimulated PBMCs from 104 healthy individuals from the Northern Netherlands, a dataset measured using version 2 chemistry (hereafter called the Oelen v2 dataset) and one measured using version 3 chemistry (called the Oelen v3 dataset), 2) a dataset from the van der Wijst study (10) that collected unstimulated PBMCs from 45 healthy individuals from the Northern Netherlands measured using version 2 chemistry (called the ‘van der Wijst’ dataset) and two datasets from the van Blokland study (19) that collected unstimulated PBMCs from 38 individuals 6–8 weeks after having a heart attack, one dataset measured using version 2 chemistry (called the van Blokland v2 dataset) and one measured using version 3 chemistry (called the van Blokland v3 dataset) (Figure 1a, Supplementary Table 1).

**Figure 1.**
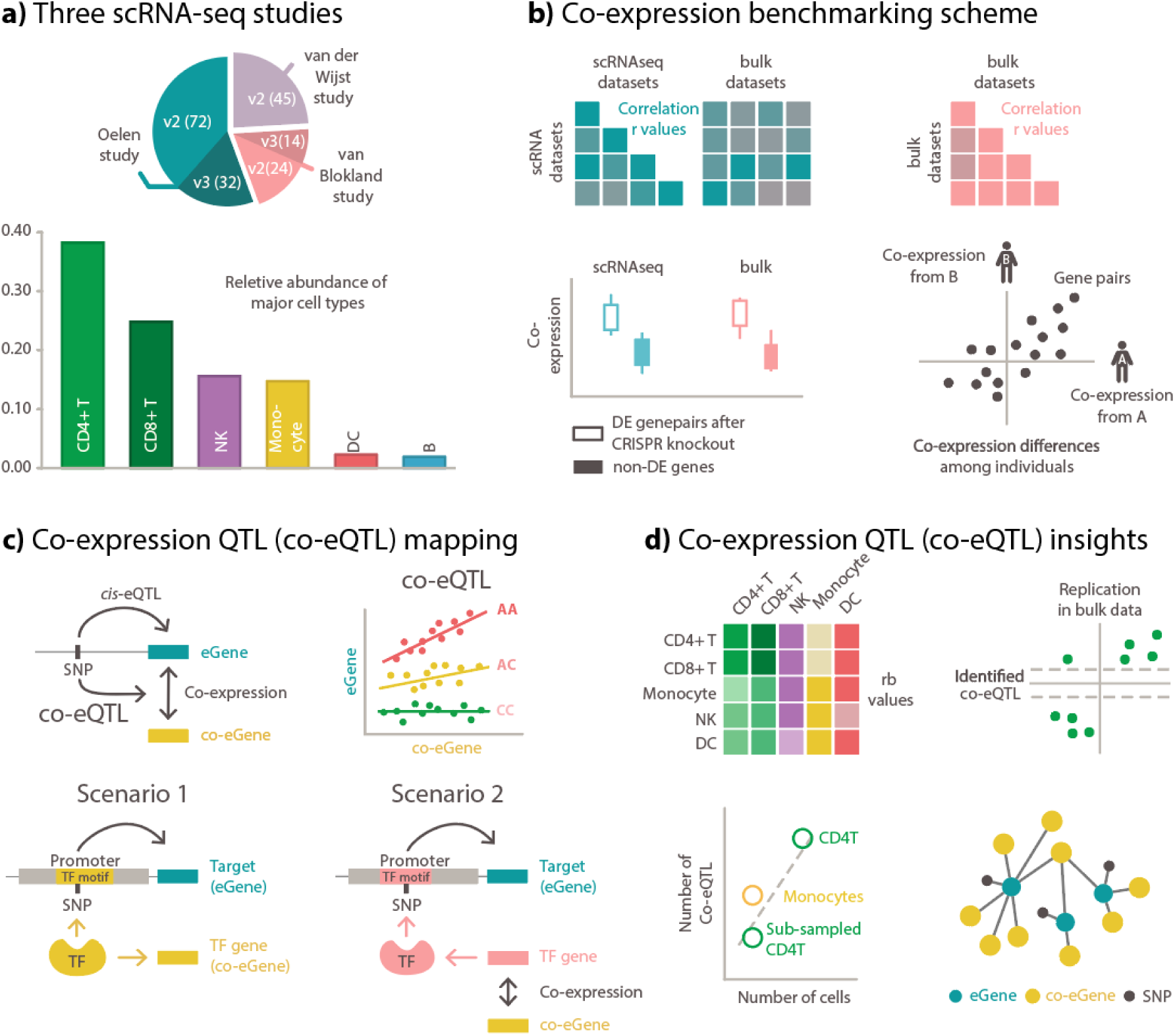
Study overview. **a)** Overview of the three PBMC scRNA-seq studies used in our study. The studies, the version of the used chemistry for data generation (version 2, referred as v2, and version 3, referred as v3), number of individuals involved (indicated as the number in the parenthesis), and relative composition of the major blood cell types used in this study. **b)** Co-expression benchmarking scheme. We first benchmarked co-expression patterns among the three scRNA-seq studies and compared them to co-expression patterns in different bulk datasets. As an additional external validation, we benchmarked both the scRNA-seq and bulk co-expression patterns with a CRISPR knockout dataset. After benchmarking, we evaluated differences in co-expression patterns among cell types and among individuals within a cell type. **c)** Co-expression QTL (co-eQTL) mapping. Building on the benchmarked co-expression pattern, we developed a novel strategy to identify co-eQTLs (genetic variants changing co-expression). Part of the strategy is a strict filtering of tested SNP–eGene–co-eGene triplets, where the SNP is required to be an eQTL for one of the genes and the genes show significant correlation in at least a certain number of individuals. **d)** Co-expression QTL (co-eQTL) insights. co-eQTL mapping was conducted for each major blood cell type, then replicated in a bulk dataset. To evaluate technical influences, we assessed the impact of cell number, number of tests and the number of individuals on the number of significant co-eQTLs. Lastly, we interpreted the biological relevance of the co-eQTLs and reconstructed the gene regulatory network using identified co-eQTLs.

We focused on the six major cell types (B cells, CD4+ T cells, CD8+ T cells, dendritic cells (DCs), monocytes and natural killer (NK) cells), of which CD4+ T cells, CD8+ T cells and monocytes were the most frequent cell types (**Supplementary Figure 1**). We compared commonly used measures of correlation and those previously reported to be particularly suitable for capturing co-expression in scRNA-seq data, including rho proportionality (16), Spearman correlation and GRNBoost2 (20), and tested complementary strategies such as MetaCell (17). We validated that the co-expression patterns from our single-cell dataset are enriched for actual gene regulatory relationships by benchmarking the concordance of the co-expression patterns across the three single-cell studies (10,11,19) and three cell-type-specific or whole-blood bulk RNA-seq datasets (2,21,22) (**Figure 1b**). Furthermore, we validated identified connections with a CRISPR dataset (23).

Next, we evaluated the concordance of the co-expression networks between the major blood cell types and between different individuals within each cell type (**Figure 1b**). To identify the genetic contribution to such common and cell-type-specific effects, we performed a constrained co-eQTL meta-analysis. For this, we filtered SNPs that exhibit an eQTL effect (with the corresponding gene referred to as an eGene below) and tested all genes with sufficient co-expression strength with the eGene (called co-eGenes below) among different individuals (**Figure 1c**).

For the co-eQTL interpretation, we considered different scenarios that can lead to detection of co-eQTL. One is that the genetic variant changes the binding affinity of a TF and thus the regulation of its target gene, which would cause a co-eQTL between the variant, the TF and the target gene (**Figure 1c**). However, a co-eQTL will also occur for all genes in strong correlation with this TF (**Figure 1c**), so we tried to identify directly interacting TFs via additional annotations and enrichment analyses. Other scenarios include genetic variants that change the structure of the TF and thereby its binding affinity and genetic variants that affect sub-cell-type composition and thus the correlation pattern of sub-cell-type-specific genes.

We then replicated the identified co-eQTLs in a large bulk study (2), explored technical factors influencing the identification of co-eQTLs (sample size, number of cells, different filtering approaches) and biologically interpreted several examples of co-eQTLs (**Figure 1d**).

### Correlation validation

Co-expression correlations can be assessed using various dependency measures. A recent benchmark study (15) reported that the proportionality measure from the propr package (16) outperforms several other methods in the identification of functional, coherent biological clusters. We observed high correlations between rho proportionality and Spearman correlations (r = 0.68) for genes expressed in > 5% of the cells (**Supplementary Figure 2a**), but for genes expressed in fewer cells, rho proportionality gave arbitrarily high values while the Spearman correlation remains near zero (**Supplementary Figure 2b**). The reason for the stark differences for very lowly expressed genes is probably that rho proportionality changes zero values to the next lowest value of the gene pair, which may result in false positive associations (i.e. very high rho values) for lowly expressed gene pairs. Another drawback of rho proportionality is the high computational demand (24), which makes it challenging to evaluate all gene pairs. As the differences between Spearman correlation and rho proportionality are very small for highly expressed genes and Spearman correlation calculation is far more efficient and handles zero values better, we chose to use Spearman correlation over rho proportionality.

We also tested other approaches, including GRNBoost2 (20), grouping cells into *MetaCells* (17) before calculation of Spearman correlation, and testing pseudotime ordering (25) and RNA velocity (26), but these did not yield more reliable results than Spearman correlation (**Supplementary Figures 3,4,5; Supplementary Text**). We therefore selected Spearman correlation to measure the co-expression patterns in scRNA-seq data for its robustness and simple interpretability. However, although we determined that Spearman correlation was optimal for the single-cell PBMC datasets that we studied, we cannot exclude that the other methods might be optimal for other single-cell datasets.

We then evaluated whether the co-expression patterns obtained from scRNA-seq data are robust and reproducible across different single-cell datasets and whether they reflect functional interactions among genes. Benchmarking the co-expression patterns obtained from scRNA-seq data is difficult because, to our knowledge, there is no clear gold-standard dataset of known functional gene–gene interactions for different cell types. As an alternative approach to assess the reliability of the identified co-expression relationships, we compared to what extent we could replicate the co-expression patterns found in one dataset in another dataset.

We first compared the cell-type-specific co-expression patterns among the five scRNA-seq datasets in our study (10,11,19). For this, we inferred the co-expression strength using Spearman correlation for each gene pair in each dataset and cell type, where gene pairs were only considered when both genes were expressed in at least 50% of the cells. We summarized the concordance between datasets by calculating the Pearson correlation on the gene pair correlation values. Overall, there was high concordance across all cell types (median r = 0.80 across all cell types). CD4+ T cells, the most abundant cell type in our dataset, had a high correlation across the different 10X chemistries and datasets, with values ranging from 0.67 to 0.86 and a median of 0.81 (**Figure 2a**). For CD8+ T cells and NK cells, we observed a comparably high correlation (CD8+ T cells median r = 0.86, NK cells median r = 0.80), while the correlation was slightly lower for the other cell types (monocytes median r = 0.69, B cells median r = 0.70, DCs median r = 0.71) (**Supplementary Figure 6**). The number of genes expressed in 50% of the cells varied between dataset and chemistry, so it was not always possible to test the same set of genes. In general, this filtering strategy is quite stringent, yielding a limited number of tested genes (at most 766 genes for the Oelen v3 dataset in CD4+ T cells, **Figure 2a**), which ensured a high-quality gene set to test due to the sparse single-cell data. A detailed evaluation of the expression cutoff follows in the next sections.

**Figure 2.**
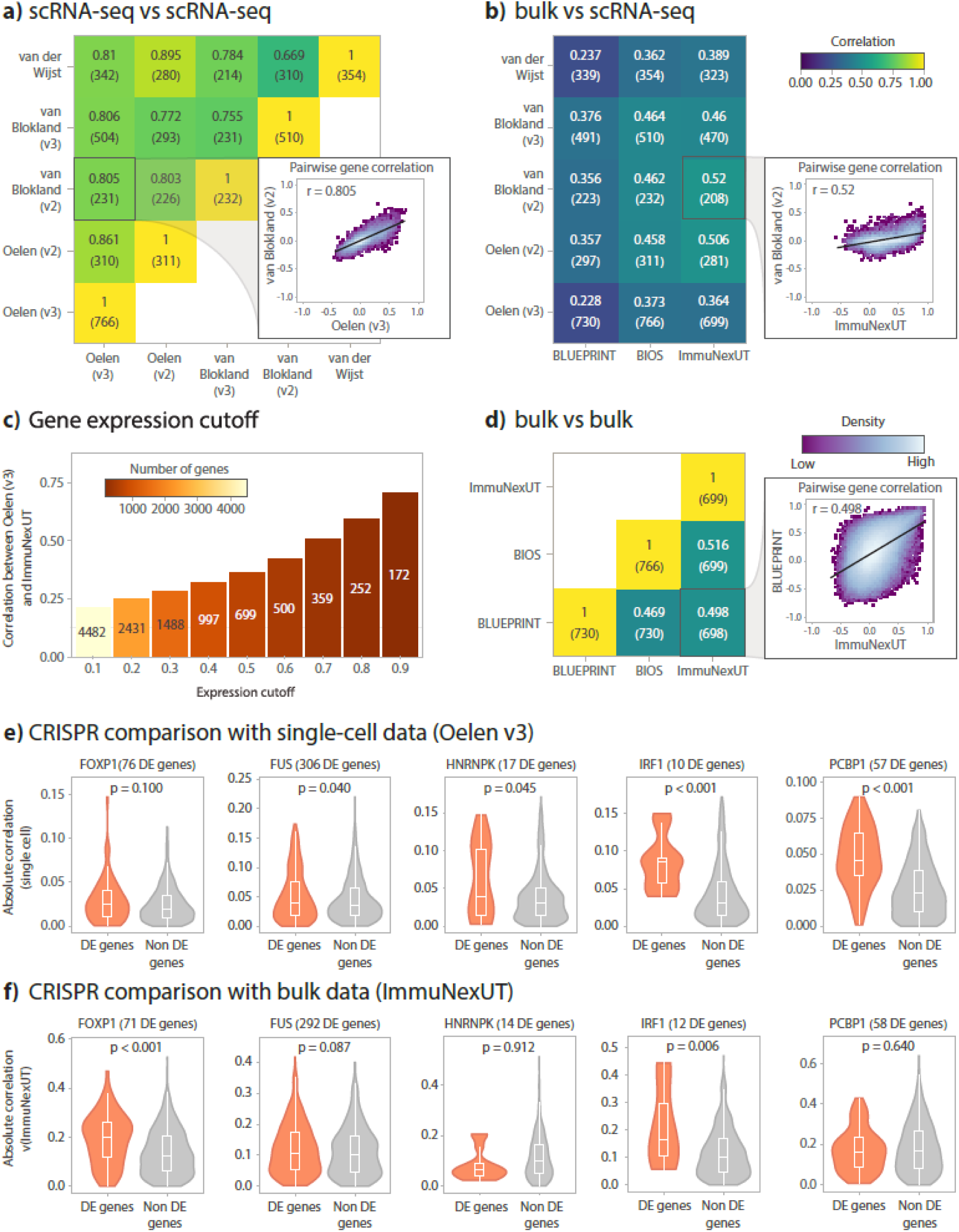
Evaluation of correlation metrics. **a)** Comparison of the co-expression profiles among the different single-cell datasets in this study. Spearman correlation of the Oelen v2 and v3 datasets, the van Blokland v2 and v3 datasets and the van der Wijst dataset were compared with each other, always taking the CD4+ T cells and genes expressed in at least 50% of the cells in the corresponding datasets. The number of genes tested is shown in parentheses below the exact Spearman correlation value. **b)** Comparison of the co-expression profiles between the single-cell datasets and with the bulk RNA-seq datasets from BLUEPRINT, ImmuNexUT (both measuring FACS-sorted naive CD4+ T cells) and BIOS (whole blood). Again, we only assessed genes expressed in at least 50% of the cells for the single-cell dataset (number of tested genes shown in parentheses below the Spearman correlation value). **c)** Relationship between the co-expression similarity between the ImmuNexUT naive CD4+ T cells and Oelen v3 dataset CD4+ T cells and increasing gene expression cutoffs (the ratio of cells with non-zero expression for a given gene). The number of genes tested are indicated by color scale and the numbers in the bar plot. **d)** Comparison of the co-expression profiles between the bulk RNA-seq datasets, taking the same gene subset as in **a)** and **b)**. The number of tested genes is shown in parentheses below the exact Spearman correlation value. **e)** Enrichment of correlated genes in scRNA-seq (Oelen v3 dataset) among associated genes identified by CRISPR knockout. For the enrichment, genes differentially expressed after knockout of FOXP1, FUS, HNRNPK, IRF1 and PCBP1 were identified and tested for enrichment. P-values in the plot show the significance level of the Wilcoxon rank-sum test. **f)** Enrichment of correlated genes in bulk RNA-seq (ImmuNexUT) among associated genes identified by CRISPR knockout. See **e)** and **Methods** for further details.

Next, we compared the co-expression patterns from the single-cell datasets to three different bulk datasets from BLUEPRINT (21), ImmuNexUT (22) and the BIOS Consortium (2). The BLUEPRINT dataset contains fluorescence-activated cell sorting (FACS)-sorted expression data from naive CD4+ T cells and classical monocytes for up to 197 individuals. The ImmuNexUT study collected gene expression data from 337 patients for 28 FACS-sorted immune cell subsets. The BIOS dataset contains whole-blood expression data from 3,198 individuals. Notably, the co-expression correlation between the single-cell and bulk-based datasets (**Figure 2b**) was much lower than those between the single-cell datasets alone (**Figure 2a**).

Comparing our single-cell data with ImmuNexUT, the only dataset with cell-type-specific expression for all evaluated cell types, CD8+ T cells showed the highest correlation (median r = 0.570) and monocytes (median r = 0.395) and DCs (median r = 0.259) showed the lowest correlations (**Figure 2b**, **Supplementary Figure 7**). The correlations from BLUEPRINT were slightly lower but in the same range (CD4+ T cells median r = 0.356, monocytes median r = 0.339) (**Figure 2b**, **Supplementary Figure 7**). Finally, we observed that the whole blood bulk data from the BIOS dataset correlated reasonably with the different single-cell cell types (median r between 0.265 and 0.458 across cell types; **Figure 2b**, **Supplementary Figure 7**).

We studied this seemingly low correlation between bulk and single-cell data, and identified multiple factors that play a role. One is the sparseness of the single-cell data, which could introduce noise and therefore lead to less stable co-expression values. To test this, we correlated the co-expression from the Oelen v3 dataset with that from ImmuNexUT using varying expression cutoffs based on the number of cells expressing a gene (**Figure 2c**). Indeed, the sparseness of the single-cell data affects the correlation. We observed increased concordance with increasing gene expression levels: the correlation increased from r = 0.21 for an expression cutoff of 10% to r = 0.71 at a cutoff of 90%. However, the number of genes that can be tested dropped from 4,482 at an expression cutoff of 10% to 172 at a cutoff of 90%. The same trends were observable when comparing the Oelen v3 dataset with the BLUEPRINT dataset for different cutoffs (**Supplementary Figure 8**). For this reason, we chose a cutoff of 50% as a trade-off between both extremes in our benchmarking study (**Figure 2a,b,e,f**).

Other aspects that may affect correlations between genes are the difference in resolution and potential biases introduced by acquiring cell-type-specific data, such as the gene expression changes induced by FACS and the technical complications of deconvoluting cell types. Furthermore, the validity of bulk-based correlations is affected by the possibility of Simpson’s paradox occurring. Simpson’s paradox describes the incorrect introduction or removal of correlations by averaging expression levels. This can potentially occur in bulk datasets, whereas single-cell data can accurately identify the co-expression value since we can calculate co-expression values per cell type and per individual (**Supplementary Figure 9a**). To estimate the effects of this phenomenon, we recalculated co-expression from the single-cell data using a bulk-like approach, compared it to the normal single-cell co-expression values and observed several examples of highly expressed genes in which Simpson’s paradox occurs (**Supplementary Figure 9b,c**). However, taking the average gene expression over many cells also results in more robust expression estimates, which can generate less noisy co-expression estimates, especially for lowly expressed genes. For this reason, we cannot differentiate for all genes which co-expression differences between single-cell and bulk are caused by Simpson’s paradox and which are caused by noisy single-cell data.

To contextualize the correlation values between single-cell and bulk data, we also compared the bulk datasets with each other and assessed whether bulk datasets actually capture gene co-expression consistently. Surprisingly, the co-expression correlation similarity between bulk datasets was quite low (r between 0.47 and 0.52 for CD4+ T cells and between 0.35 and 0.42 for monocytes) (**Figure 2d, Supplementary Figure 10**). Given that these correlations are expected to be an upper bound when comparing bulk datasets with single-cell datasets, our observed correlations in those comparisons are very reasonable.

Given the imperfect correlation between the different bulk datasets, we used gene expression data from CRISPR-knockouts as an additional evaluation criterion. For this purpose, we benchmarked the co-expression patterns from our single-cell datasets against a CRISPR knockout scRNA-seq dataset in CD4+ T cells (23). While a unique single-guide RNA barcode reveals which gene was targeted in which cell, some cells may escape from successful CRISPR perturbation. To account for this, we used Mixscape to assign a perturbation status to each cell (27). For each knockout, we then determined other genes that were differentially expressed (DE) in successfully perturbed cells compared to wild-type cells. We then selected genes for which perturbation resulted in at least 10 DE genes and compared the correlation of these DE genes with non-DE genes using the Wilcoxon rank-sum test (see **Methods**). For four out of five gene knockouts, we observed significantly higher correlation of the knockout gene with the DE genes than with non-DE genes (p < 0.05) in the single-cell dataset (**Figure 2e**). In contrast, the bulk naive CD4+ T cell data from ImmunNexUT showed a weaker connection between correlation and DE genes, with only two out of five knockout genes having significantly higher correlation with the DE genes (p < 0.05) (**Figure 2f**).

As another line of evidence, we tested whether pairs of genes known to interact on the protein level showed higher co-expression correlation compared to other pairs of genes. Here we found that gene pairs with protein interactions listed in the STRING database (28) had a higher co-expression correlation than gene pairs not in STRING, both when using the single-cell dataset and the bulk dataset (for both Wilcoxon rank-sum test, p < 0.05, **Supplementary Figure 11**).

Overall, we have shown that single-cell data can identify true gene co-expression relationships as co-expression patterns from scRNA-seq data are highly replicable among different datasets and are supported by functional interactions among genes identified by CRISPR perturbations and the STRING database.

### Cell type and donor differences in co-expression pattern

Next, we examined cell-type-specific and individualized co-expression patterns. As expected, lymphoid cell types (B, T and NK cells, r > 0.73) were more alike with each other but they are less alike with myeloid cell types (monocytes and DCs, r > 0.45) (**Figure 3a**, **Supplementary Figure 12a**). However, myeloid cell types were not as alike to each other as lymphoid cell types. This is possibly due to the fact that DCs are one of the least abundant cell types (**Supplementary Figure 1**), which would have resulted in less accurate co-expression estimations. Overall, the correlation between different cell types within one scRNA-seq dataset (for Oelen v3 dataset median r = 0.64, **Figure 3a**) was generally lower than the correlation between different scRNA-seq datasets when studying a single cell type (median r = 0.80 across all cell types, **Figure 2a, Supplementary Figure 6**). These differences highlight cell-type-specific differences in the correlation pattern, further confirming the biological aspects captured by scRNA-seq co-expression values. We also explored the distribution of co-expression among cell types (**Figure 3b, Supplementary Figure 12b**). Typically, the correlations between gene pairs were rather low, with only a small proportion of gene pairs (median 12.4%) showing correlations above 0.1. However, we did observe cell-type-specific differences, with DCs possessing a higher proportion of co-expressed gene pairs compared to the other cell types (32.3% of gene pairs with r > 0.1).

**Figure 3.**
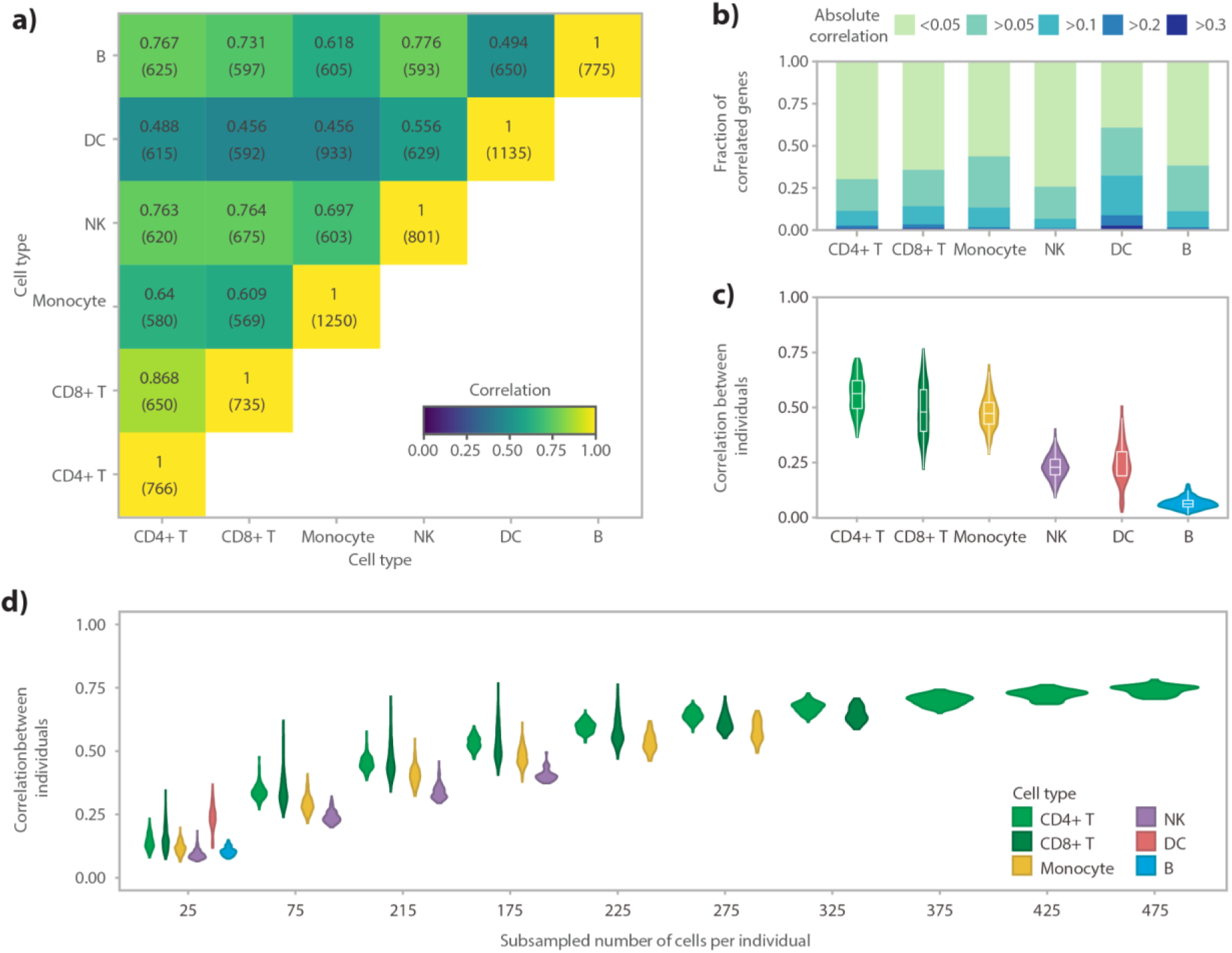
Comparison of correlation across cell types and donors. Each analysis was performed in the Oelen v3 dataset for all genes expressed in at least 50% of the cells of the respective cell type. **a)** Comparing co-expression patterns across cell types within the Oelen v3 dataset for genes expressed in 50% of the cells for both cell types in each pair-wise comparison. The number of tested genes is shown in parenthesis below the Spearman correlation value. **b)** Correlation distribution within each cell type. **c)** Correlation between different individuals within each cell type showing the distribution of all pair-wise comparisons between individuals. **d)** Relationship between the number of cells per individual and cell type and correlation between individuals separately for each cell type. In each subsampling step, we assessed all individuals who have at least this number of cells and subsampled to exactly this number (this leads to removal of some individuals for higher number of cells and thus, a direct comparison with the correlation values in **c)** is not possible).

In addition to detecting cell-type-specific associations, scRNA-seq enables direct calculation of co-expression correlations per individual as it provides many measurement points per donor. When we calculated the correlation separately for each donor and cell type, we observed overall strong correspondence of co-expression networks between different donors for the more frequent cell types (CD4+ T cells median r = 0.56, CD8+ T cells median r = 0.48, monocytes median r = 0.47) (**Figure 3c, Supplementary Figure 12c**). As a result of noisier estimates, the correlation between individuals was drastically lower for the less frequent cell types (DCs median r = 0.24, B cells median r = 0.06). Moreover, these correlations were much smaller than comparing one cell type across entire datasets (i.e. including all individuals at once), which showed correlations of at least 0.81 for CD4+ T cells, 0.64 for CD8+ T cells, 0.49 for monocytes, 0.66 for NK cells, 0.62 for B cells and 0.38 for DCs (**Figure 2a**). This decline is potentially caused by the number of cells used to calculate the correlation, which is drastically lower when comparing donors within one dataset. The number of cells could also explain the differences between the cell types. To test this, we subsampled the number of cells for each cell type and indeed observed that the correlation increased when the number of cells increased (**Figure 3d**). Apart from the number of cells, we also observed potential cell type differences. The similarities between individuals were significantly smaller in NK cells compared to monocytes and T cells, when the same number of cells was used (**Figure 3d**). We also confirmed these observations in another scRNA-seq dataset (**Supplementary Figure 12d**).

We further explored the relationship between the number of cells per individual and the correlation between individuals by fitting a logarithmic curve for the four most frequent cell types: CD4+ T cells, CD8+ T cells, monocytes and NK cells (**Supplementary Figure 13**). Each of the observed trends could be fit well with the logarithmic curve (adjusted R^2^ values between 0.86 and 0.98). We then extrapolated the trend to 1,000 cells, showing that a correlation > 0.80 would be expected for T cells and monocytes with this number of cells and a correlation of 0.65 for NK cells (**Supplementary Figure 13**). We acknowledge, however, that the exact upper bound for the correlation between donors cannot be estimated accurately with our current dataset. For example, the correlation close to 100% for CD4+ T cells and 1,500 cells is likely too high considering that donor-specific differences such as genetics and environment will remain independent of the number of cells. Nevertheless, our fits highlight the value of having measurements from many cells for accurate correlation estimates as well as cell-type-specific differences in the correlation pattern.

During this comparison, we observed a few gene pairs that showed a high variance in correlation across donors within one cell type (median fraction of gene pairs with correlation Z-score variance > 2 across cell types: 4.9% for Oelen v2 dataset and 3.3% for Oelen v3 dataset, **Supplementary Figure 14**). This high variance could, in theory, be caused by different sources, e.g. technical factors or environmental influences, but could also reflect genetic differences between individuals. Since we observed low co-expression variance between different individuals for the same cell type and similar numbers of cells (**Figure 3d**), we concluded that these differences are not likely to originate from technical factors, and thus we next looked into genetic variation as one of the other potential major influences.

### Establishing a method to identify co-expression QTLs

To assess how strongly genetic variation influences the correlation between pairs of genes, we performed a co-eQTL analysis. In contrast to classical eQTL analysis, co-eQTL analysis not only reveals the downstream target gene whose expression is affected by a genetic variant, it can also help identify the upstream regulatory factors that affect these eQTLs, as discussed in the Overview.

Compared to an eQTL analysis, a full co-eQTL analysis with all SNP–gene pair combinations would massively increase the multiple testing burden. Previously, we showed the necessity of filtering the SNP–gene pair combinations to reduce the multiple testing burden associated with a genome-wide co-eQTL analysis on all possible triplets while not missing true co-eQTLs (11). For example, in our current study, testing all pairs of genes expressed in monocytes would lead to 1.96×10^8^ tests when considering only one SNP per pair and to a very limited power to detect small effect sizes (power of 1.4% to detect a significant effect for a phenotype (here the co-expression relationship) with a heritability of 10% that is explained by a single locus, **Supplementary Figure 15**).

In this study, we aimed to define a generally applicable filtering strategy that yields a large number of highly confident co-eQTLs. First, we decided to focus on *cis*-eQTL SNPs and genes because we expect a SNP influencing the co-expression of two genes to also influence the expression of one of the genes directly (a strategy we applied successfully before in (10, 11)). To identify these *cis-*eQTLs, we first performed a *cis*-eQTL meta-analysis across four of the five scRNA-seq datasets. We excluded the van Blokland v3 dataset from this eQTL analysis and all subsequent analyses because the small sample size (N = 14) provided very few variants above the minor allele frequency (MAF) cutoff (>10%), which made it unsuitable for this meta-analysis. To reduce the multiple testing burden and maximize the number of *cis*-eQTLs detected given the relatively low number of individuals (N = 173) used for the eQTL mapping, we confined ourselves to 16,987 lead *cis*-eQTLs previously identified in a large (N = 31,684) bulk blood eQTL study (2). Depending on cell type, we identified between 917 (for CD4+T cells) and 51 (for B cells) eQTLs (FDR<0.05; **Supplementary Table 2, 3**).

As filtering for the eQTL effects still resulted in a large number of tests (e.g. for CD4+ T cells, n = 12,137,281, **Supplementary Table 4**) and consequently a large multiple testing burden, we imposed additional filtering on the co-eGenes to study. Here, we used a filtering strategy based on the co-expression significance, selecting co-eGenes for which we observed a significant (nominal p ≤ 0.05) correlation with the eGene in at least 10% of the individuals (**Methods**). We assumed this captures genuine co-expression effects that are present in at least one of the genotype groups (i.e. homozygous reference/heterozygous/homozygous alternative allele). Note that the filtering strategy we used here is less stringent than the cutoff used in the co-expression benchmarking analyses (**Figure 2,3; Methods**). This is because the two analyses have very different goals, while the benchmarking was more technical in nature, we aimed to uncover new biology in the co-eQTL analyses. Thus we used a less stringent selection in the co-eQTL analysis to ensure that we did not miss out on detecting relevant biological processes underlying gene regulation.

An additional challenge is the large number of missing co-expression values for gene pairs within individuals. This is introduced by the sparsity of the scRNA-seq data: correlation is missing when the expression of one gene is zero in all cells of an individual. We argue that these missing co-expression values may not reflect true null correlations between gene pairs because zero values in single-cell data can also be caused by lowly expressed genes not being quantified accurately. As we observed that replacing missing values with 0 can lead to spurious co-eQTL results (**Supplementary Figure 16**), we removed such missing correlations when calculating co-eQTL correlations rather than imputing them with 0 correlation values.

Finally, we applied a customized permutation strategy for each gene pair. Since common upstream regulators might lead to co-expression of many co-eGenes, we expect correlated test statistics among the family of tests carried out for each SNP– eGene pair. Therefore, we applied a customized permutation strategy per SNP–eGene pair and an adapted multiple testing correction strategy based on fastQTL (29, 30) (see **Methods** for details).

With our co-eQTL mapping strategy, we conducted a meta-analysis with four of the five single-cell datasets (Oelen v2 and v3, van Blokland v2 and the van der Wijst dataset). This identified cell-type-specific co-eQTLs for 72 independent SNPs, affecting 946 unique gene pairs in total (**Supplementary Table 5, Supplementary Table 6**). We identified the maximum number of 500 co-eQTLs in CD4+ T cells, comprising 30 SNPs, 500 gene pairs and 420 unique genes. We identified the minimum number of 35 co-eQTLs in B cells, comprising 1 SNP, 35 gene pairs and 36 unique genes.

We first examined the cell-type-specificity of these co-eQTLs. This analysis is limited by the fact that, due to our filtering strategy, we used a different set of cell-type-specific eQTLs and tested a different set of co-eGenes. Consequently, this resulted in very different sets of tested triplets for biologically different cell types, which could explain the small overlap of significant co-eQTLs between cell types (**Supplementary Figure 17, 18a; Supplementary Table 6**). Therefore, to give a complete picture of the cell-type-specificity of co-eQTLs, we replicated co-eQTLs from each cell type in all other cell types and quantified this with two different measures: 1) the ratio of co-eQTLs that could be tested in the replication cell type (**Supplementary Figure 18a**) and 2) the r_b_ concordance measure (31), which reflects the correlation of the effect sizes for the co-eQTLs that were tested in the replication cell type (**Figure 4a, Supplementary Figure 18b,** details in **Methods**). Consistent with the co-eQTL overlap results, the ratio of tested co-eQTLs are generally small, ranging between 5% to 97% (**Supplementary Figure 18a**). However, for the SNP–eGene–co-eGene triplets that were tested in the replication cell type, their effect sizes and directions were generally highly concordant, with a median r_b_ value of 0.85 (**Figure 4a, Supplementary Figure 18b, Supplementary Table 7**). The highest r_b_ were observed between CD4+ T cells and CD8+ T cells (0.97 for co-eQTLs identified in CD4+ T cells replicated in CD8+ T cells, for co-eQTLs identified in CD8+ T cells replicated in CD4+ T cells).

**Figure 4.**
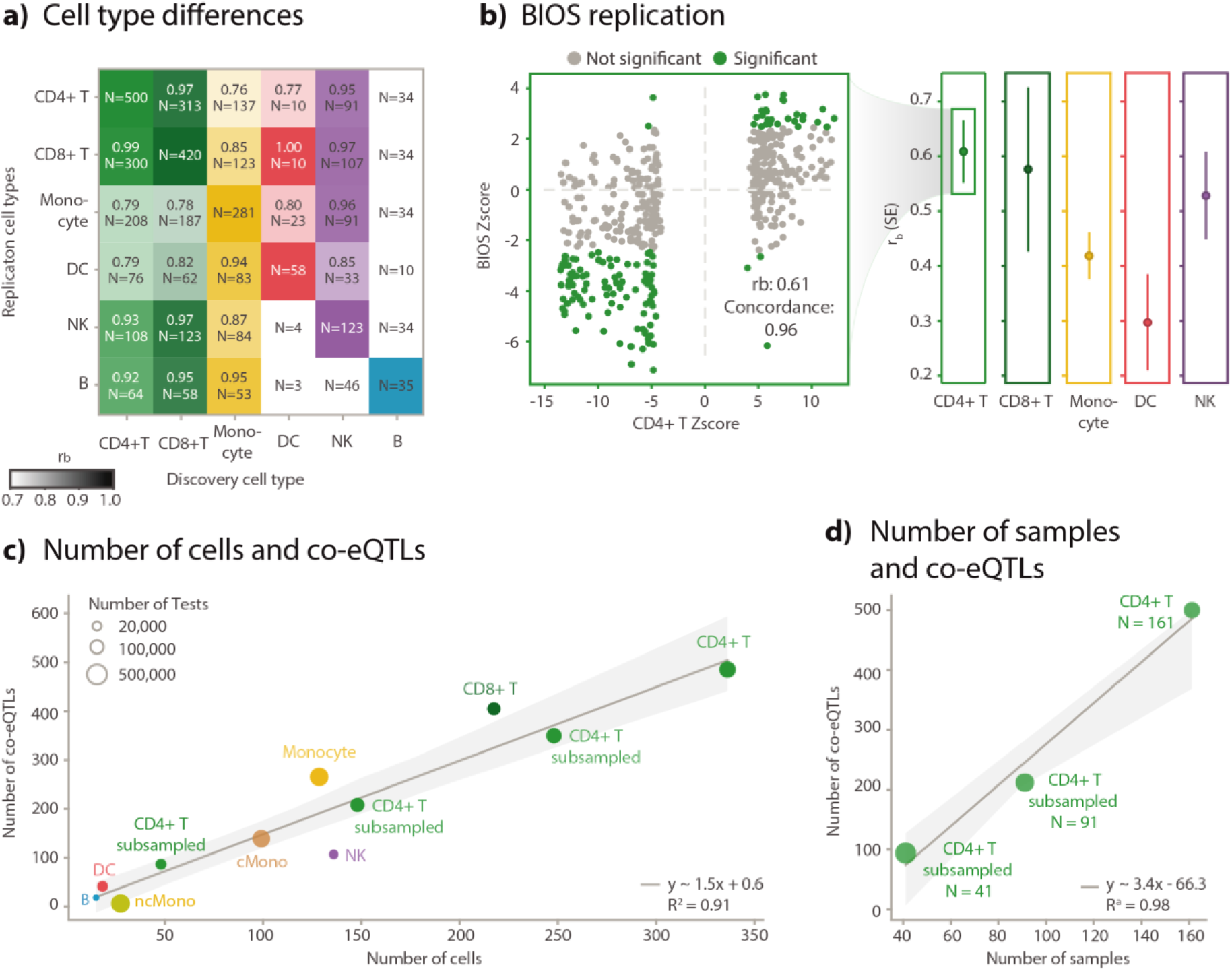
General characteristics of identified co-eQTLs. **a)** Replication of discovered co-eQTLs across the major cell types. Correlation of the effect sizes in replications among different cell types, measured by r_b_ value. Text inside each block indicates the r_b_ value, standard error and number of replicated co-eQTLs. Color intensity indicates r_b_ value. **b)** Replication in BIOS dataset for different cell types, indicated by the r_b_ values. Scatter plot shows the detailed Z-score comparison between the co-eQTL meta-analysis and the Z-score from the BIOS replication for CD4+ T cells. **c)** Number of significant co-eQTLs for varying cell numbers. Dot color indicates the cell type, as indicated in the text next to each dot. “cMono” means classical monocytes. “ncMono” means non-classical monocytes. “CD4+ T Subsampled cells” means that this analysis was done for CD4+ T cells, but for every individual we randomly downsampled cells to the desired cell number as indicated in the x-axis. **d**) Number of significant co-eQTLs for varying sample numbers. “CD4+ T Subsampled Individuals” indicates that this analysis was done for CD4+ T cells, but we randomly subsampled for the individuals.

To validate our co-eQTL results, we first examined the effect sizes and directions among the datasets used in the meta-analysis and observed high correlations (**Supplementary Figure 19**). Next, we replicated them in the BIOS bulk whole blood dataset (N = 2,491 excluding common individuals, see **Methods**) (2), using the ratio of tested co-eQTLs and r_b_ value (see **Methods**). For this replication, we used a linear regression model with an interaction term to model the associations between the expression level of eGenes and the product of genotype and the expression level of co-eGenes (see **Methods** for detailed explanation), as we have done before (8). We tested all identified co-eQTLs in the BIOS data. and their effect sizes and directions showed r_b_ values between 0.30 to 0.61 (**Figure 4b, Supplementary Table 8, 9**), with the highest concordance achieved for CD4+ T cells, with an r_b_ value of 0.61 (SE = 0.06).

After we established a baseline for the number of co-eQTLs identified and their replication rates, we used this to evaluate various technical factors such as filtering strategy, sub-cell-type composition, sample size and cell number. We first compared the analysis to a set of co-eQTLs identified when omitting the filtering step for significantly correlated gene pairs, which increased the number of tests (**Supplementary Table 4**). While this led to detection of an increased number of co-eQTLs for the more abundant cell types (CD4+ T, CD8+ T, monocytes and NK cells) and a decreased number of co-eQTLs for less abundant cell types (B cells and DCs) (**Supplementary Tables 4, 10**), we also observed a general decrease in concordance among cell types compared to the co-eQTLs obtained with the filtering strategy (**Supplementary Figure 18, 20; Supplementary Table 11**). We then repeated the BIOS replication procedures for co-eQTLs found without the filtering strategy and observed a decrease in effect concordance compared to the set of co-eQTLs identified with the filtering strategy (**Supplementary Figures 21-23; Supplementary Tables 12, 13**), indicating that the filtering increases the robustness of the co-eQTLs.

We additionally explored the correlation mean and variance, as well as the non-zero ratio for co-eQTLs compared to non-significant triplets, in the scenarios with and without additional filtering (**Supplementary Figure 24**). Here we observed that significant co-eQTLs show both a higher co-expression correlation mean and variance and a higher non-zero ratio for their expression (**Supplementary Figure 24**) compared to non-significant triplets. This is to be expected as gene pairs with a high average co-expression correlation more likely reflect true biological associations and gene pairs with a high correlation variance likely reflect true co-expression network polymorphisms. This trend is also much clearer for the filtered set compared to the non-filtered set (**Supplementary Figure 24**), suggesting that alternative preselection strategies could be envisioned that are based on specific expression values or co-expression correlation variance thresholds.

Sub-cell-type composition is a potential confounder that might introduce false positive co-eQTLs, similar to cell type-composition in bulk studies (32). If a genetic variant is associated with sub-cell-type composition, co-eQTLs with sub-cell-type-specific genes might be identified even when there is no direct association between the SNP and the co-expression. To assess this, we analyzed co-eQTLs found among classical monocytes, non-classical monocytes and the whole set of all monocytes. Here we found that co-eQTL effect sizes are highly concordant (r_b_ ≥ 0.9) (**Supplementary Figure 25**) for co-eQTLs tested in one of the subtypes and in the major cell type (>82% of co-eQTL identified in monocytes were tested in both classical monocytes and non-classical monocytes). This suggests that the co-eQTLs are not generally driven by sub-cell-type composition, although individual co-eQTLs could still be caused by sub-cell-type differences.

To highlight how future co-eQTL analyses can benefit from the expected expansion of population-based scRNA-seq datasets with available genotype data, we determined how the number of identified co-eQTLs is related to the number of individuals and cells per individual. To test the influence of the number of cells, we randomly subsampled the CD4+ T cells and monocytes per individual and repeated the co-eQTL mapping pipeline (**Figure 4c**). For the influence of the number of individuals, we randomly subsampled the individuals for CD4+ T cells (**Figure 4d**). We observed that the number of co-eQTLs is linearly and positively correlated with both the number of cells and the number of individuals, although the number of individuals had a stronger effect than the number of cells (**Figure 4c,d; Supplementary Table 5**).

### Annotating identified co-expression QTLs

After we successfully validated the identified co-eQTLs by exploring different technical aspects and replicating them in the BIOS dataset (2), we examined to what extent the co-eQTLs could provide interesting biological insights into genetic regulation, which could be relevant for the interpretation of disease variants. As discussed in the Overview, we hypothesize that among the co-eGenes identified for each SNP–eGene pair there are direct regulator genes or genes co-expressed with the direct regulators for the eGene. Even if the direct upstream regulatory factor was not evaluated in the co-eQTL analysis, due to the limited capturing efficiency of the single-cell data, the biological function of the co-eQTLs could still be inferred by the other co-eGenes in strong co-expression with the unknown upstream regulator as they presumably share the same biological function and potentially also a common role in disease. To assess these hypotheses, we combined different lines of evidence: functional enrichment based on gene ontology (GO) terms, enrichment of TF binding sites and enrichment of GWAS annotations.

Each enrichment analysis was run separately per cell type and for all co-eGenes associated with the same SNP–eGene pair (see **Methods** for details). To increase the power of enrichment analyses, we restricted ourselves to SNP–eGenes pairs with at least five co-eGenes, which covered 25% of SNP–eGenes pairs in at least one cell type (19 out of 76 unique SNP–eGene pairs). GO enrichment analysis revealed shared functional pathways for the co-eGenes. For 18 of the 19 SNP–eGene pairs, we found enrichment among the associated co-eGenes for at least one GO term (**Supplementary Table 14**). Moreover, we assessed potential common TFs regulating the shared function of these co-eGenes using ChIP-seq data processed by ReMap 2022 (33) and found enrichment of TF binding sites among the co-eGenes for 7 of the 19 SNP–eGene pairs (**Supplementary Table 15**). For four of the SNP–eGene pairs, the co-eQTL SNP itself or a SNP in high linkage disequilibrium (LD) (R^2^ ≥ 0.9) lay in the binding region of the enriched TFs (**Supplementary Table 15**), making these likely candidates for the direct regulator.

We also explored whether co-eQTLs and the respective sets of co-eGenes could enhance our understanding of disease-associated variants. For this, we annotated co-eQTL SNPs with GWAS loci, identifying approximately half the SNPs to be in high LD (R^2^ ≥ 0.8) with a GWAS locus (41 out of 72 SNPs, **Supplementary Table 16**). To assess if sets of co-eGenes for a specific SNP-eGene share a common role in disease, we explored if the co-eGenes show higher gene level trait association for GWAS traits that are also associated with the respective co-eQTL SNP. We identified overlapping GWAS traits for two co-eQTL SNPs and their co-eGenes for at least one GWAS trait and cell type, with many of the traits covering blood cell counts and immune-mediated diseases (GWAS SNP p-value < 5×10^−8^, FDR <0.05, **Supplementary Table 17**), further strengthening the biological connection of the co-eGenes with the eQTL.

Furthermore, we observed that the direction of effect of the co-eQTLs can be helpful in grouping genes sharing the same functions. For this, we compared the direction of effect of the co-eQTL with the direction of the associated eQTL, choosing the same reference allele in both cases. If the direction matched, we classified it as concordant. In these co-eQTLs, increasing expression of the eGene led to increasing co-expression. If the directions did not match, we said the direction of the co-eQTL is discordant. Between 37% and 97% of the co-eQTLs showed a concordant direction of effect across cell types (**Supplementary Figure 26**), but the majority of co-eGenes were associated with rs1131017–*RPS26* and thus the observed distributions are probably not generalizable for future larger studies that identify more co-eQTL.

In the following section, we highlight some examples of how these co-eQTL can help to better understand the molecular functional consequences of genetic variants associated to disease.

When grouping co-eQTLs based on their associated eQTL, eQTL rs1131017–*RPS26* had the most significantly associated co-eGenes in all cell types except for DCs (between 372 co-eGenes for CD4+ T cells and 35 for B cells) (**Figure 5a,b,c,d**). *RPS26*, encoding a ribosomal protein, showed strong correlation with other ribosomal proteins, and we had previously reported a few *RPS26* co-eQTLs in CD4+ T cells (10) and monocytes (11). Our new methodology and the larger sample size in the current study allowed us to now compare the genes part of the rs1131017–*RPS26* co-eQTLs across cell types.

**Figure 5.**
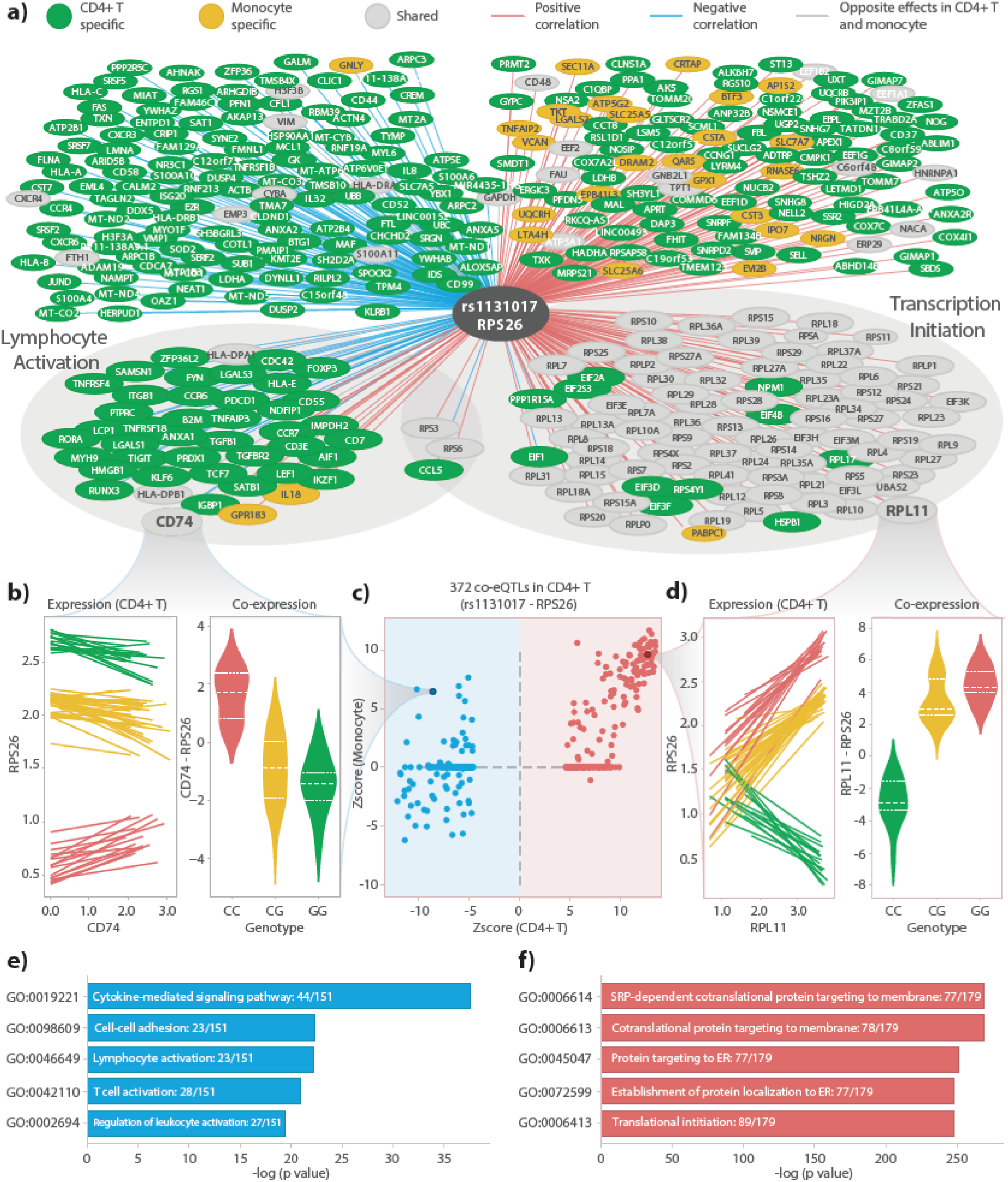
Annotation of co-eQTLs. **a**) Union network constructed with co-eQTLs found in CD4+ T cells or monocytes that are associated with the SNP–eGene: rs1131017–RPS26. The two circled clusters contain co-eGenes that are in those corresponding GO terms. **b**) Example of one co-eQTL: rs1131017–RPS26-CD74. Left plot indicates the co-expression patterns from all individuals in the Oelen v3 dataset. Each regression line was fitted with expression data from one individual. Right plot indicates the co-expression values from the three genotype groups. **c**) Comparison between z-scores from monocytes and z-scores from CD4+ T cells. Red dots indicate positive co-eQTLs from CD4+ T cells. Blue dots indicate negative co-eQTLs from CD4+ T cells. **d**) Example of one co-eQTL: rs1131017–RPS26-RPL11 with the same layout as (b). **e**) GO term enrichment results for the co-eGenes in negative co-eQTLs from CD4+ T cells. **f**) GO term enrichment results for the co-eGenes in positive co-eQTLs from CD4+ T cells.

In monocytes, NK cells and B cells, nearly all the associated genes showed a positive direction of effect, concordant with the eQTL direction (95% of all co-eGenes for monocytes, 90% for NK cells and 97% for B cells), while in CD4+ T cells and CD8+ T cells, several genes showed a negative direction of effect, discordant with the eQTL direction (46% of all co-eGenes for CD4+ T cells and 43% for CD8+ T cells).

The positively associated genes replicated well across all cell types (**Figure 5c, Supplementary Figure 27**) and were enriched for functions associated with translation (**Figure 5e**), which is consistent with the fact that many co-eGenes were ribosomal proteins from both the large and the small subunit (for CD4+ T cells: 46 of 47 tested RPL genes and all 31 tested RPS genes were associated). In contrast, the negatively associated genes only replicated well between CD4+ T cells and CD8+ T cells (**Supplementary Figure 27; Figure 5c,d**), despite the fact that these genes were well expressed in the other cell types. This negatively associated set of genes showed enrichment in functions associated with immune response and T cell activation (**Figure 5f**).

TF enrichment analysis identified six TFs—*RBM39, TCF7, LEF1, KLF6, CD74* and *MAF*—whose binding sites were enriched in the promoter region of the rs1131017– *RPS26* co-eGenes, that had a binding site overlapping with rs1131017 and that were among the rs1131017–*RPS26* co-eGenes themselves (**Supplementary Table 15**). This led us to the assumption that one or more of these TFs represent the direct regulators of the eQTL, as described in the Overview (**Figure 1c, Scenario 1**). Five of the TFs (*TCF7, LEF1, KLF6, CD74* and *MAF*) are also connected with lymphocyte activity (the first four based on GO annotations, *MAF* based on a recent study (34)), further strengthening the link with T cell activation. Of these, *MAF* and *CD74* were specifically enriched not only among all co-eGenes but additionally among co-eGenes with a negative effect direction (**Supplementary Table 15**).

GWAS enrichment analysis showed enrichment for several different blood cell counts, but only in CD4+ T cells and CD8+ T cells we also observed enrichment for immune-mediated disease (rheumatoid arthritis, Crohn’s disease (CD), multiple sclerosis (MS) and hay fever), further connecting the co-eGenes especially with T cells (**Supplementary Table 17**). Interestingly, several studies have highlighted a connection of RPS26 with T cell activation and survival (35, 36), and the associated co-eQTL SNP rs1131017 is associated with the enriched immune-mediated diseases (rheumatoid arthritis, CD, MS, hay fever), and additionally associated with type 1 diabetes (T1D) and with other autoimmune traits (37).

We examined whether the large number of co-eQTLs for rs1131017 were confounded by sub-cell-types in CD4+ T cells. We cannot exclude the possibility that this variant showed this effect in CD4+ and CD8+ T cells by specifically affecting the amount of circulating CD4+ or CD8+ sub-cell types whose marker genes would subsequently show up as co-eQTLs in our analysis, where we have not distinguished between sub-cell types. To test whether this is a possibility, we associated SNP rs1131017 and the ratio between CD4+/CD8+ TEM cells and CD4+/CD8+ naive T cells, but we did not see a significant correlation (**Supplementary Figure 28**). Together, these results suggest that RPS26 plays a dual-function role, both in general transcription and specifically in lymphocytes in T cell activation. This points to a potential working mechanism in the role of rs1131017 in the manifestation of autoimmune diseases.

Another set of promising co-eQTLs are those associated with rs7806458–*TMEM176A* in monocytes (11 co-eGenes) and rs7806458–*TMEM176B* in monocytes (6 co-eGenes) and DCs (1 co-eGene) as they connect the co-eQTL SNP rs7806458 that has been associated with MS (38) with blood coagulation. This is interesting as this disease has previously been connected to disturbances in blood coagulation (39). The relevance of the co-eGenes to MS is supported by two lines of evidence. Firstly, GO enrichment suggested that the six co-eGenes associated with rs7806458–*TMEM176B* in monocytes are enriched for complement component C3b binding (**Supplementary Table 14**), which is closely related to the blood coagulation system (40). When looking closely at the exact gene functions, we found three genes (*ITGB1, FCN1* and *CFP*) that contribute to the local production of complement (41). Secondly, GWAS enrichment analysis showed MS enrichment for co-eGenes associated with rs7806458–*TMEM176A* in monocytes (**Supplementary Table 17**). Intriguingly, the eGene *TMEM176B* was previously found to be associated with the maturation of DCs (42), and it has been shown that white blood cells, including DCs, can act as a local source of certain complement proteins (43, 44). Though we could not identify (in)direct regulator genes for these co-eQTL in our TF enrichment analysis with the ReMap database, we argue that these co-eGenes, supported by several lines of evidence, provide valuable mechanistic insights for the MS SNP rs7806458.

For several of the other co-eQTLs, we could not provide as strong and coherent evidence for the interpretation but nevertheless found promising connections to biological functions and disease that can be explored in further studies. One is the SNP–eGene pair rs9271520–*HLA-DQA2*. We found co-eQTL effects for it in CD4+ and CD8+ T cells, monocytes and DCs, with the number of co-eGenes ranging from 7 to 17. Interestingly, rs9271520 is in LD with several immune disease SNPs where we also found enrichment for the co-eGenes in the same GWAS traits. The most significant (sorted by GWAS SNP p-values) enriched traits include rheumatoid arthritis, MS and asthma (see **Supplementary Table 17** for full GWAS enrichment results). However, we found several other genes in the HLA region being co-eGenes associated with rs9271520–*HLA-DQA2*, and, when we removed those HLA genes, the GWAS enrichment signals disappeared. This indicateed that the enriched signal could be due to the LD structure in the HLA region and a confident mapping of the causal regulatory connections is not possible with our dataset. Other interesting co-eQTL examples and their interpretations are discussed in **Supplementary Text**.

In general, our study is still underpowered in finding a lot of associated co-eGenes (**Figure 4c,d**, **Supplementary Figure 15**). This limits the set of SNP-eGenes, for which we can perform a well powered enrichment analysis and so the biological interpretation of these co-eQTLs. One of the potentially interesting SNP-eGenes, with too few co-eGenes for the enrichment analysis, is rs393727 - *RNASET2*, which is associated with four co-eGenes (*B2M, ITGB1, ALOX5AP, CRIP1*). The SNP rs393727 is in very high LD with two previously described SNPs associated with Crohn’s disease (CD) and inflammatory bowel disease (IBD) (**Supplementary Table 16**), eGene *RNASET2* has also been previously associated with IBD (45), and among the four co-eGenes, *ITGB1* was associated with CD (46) and *CRIP1* is associated with gut immunity (47).

Intriguingly, we found a number of overlapping co-eGenes associated with different SNP–eGene pairs, indicating potential common upstream regulatory pathways. For example, all the co-eGenes positively associated with rs4147638–*SMDT1* are also found to be positively associated with rs11311017–*RPS26*, while the four co-eGenes negatively associated with rs393727–*RNASET2* are also negatively associated with rs1131017–*RPS26* (**Supplementary Figure 29**).

## Discussion

In this study, we validated the use of scRNA-seq data to identify cell-type-specific co-expression patterns and developed a novel approach to extend the discovery of co-eQTLs. Applying this to a large meta-analysis with 173 samples, we identified 72 independent SNPs leading to co-eQTLs for 946 unique gene pairs across different cell types. These co-eQTLs shed light on the biological processes upstream of individual *cis*-eQTLs, such as that seen for rs1131017, which affects *RPS26* expression levels and is associated to autoimmune diseases. We observed that this variant affects T cell activation genes, providing a potential explanation for the association of this variant to autoimmune diseases.

In this study, we used Spearman correlation to quantify the co-expression patterns from scRNA-seq data because of its straightforward interpretability, scalability, robustness against outliers and high reproducibility among different scRNA-seq and bulk RNA-seq datasets. However, we acknowledge that such correlations do not take into account the sparseness of scRNA-seq data, and it is difficult to infer direct regulator genes. This of course also depends on the quality of the single-cell data. Direct interactions can only be distinguished from indirect interactions when the direct upstream target was measured, which is currently not always the case. As other association methods (16, 20) that were top-performing in independent benchmarking studies (15, 18) did not perform better in our validation and a reliable temporal ordering of the cells (25, 26) was not possible in our dataset, we applied Spearman correlation as a solid basis for the co-eQTL analysis. Future work may find that other association measures are equally suitable or more suitable, and this may potentially depend on the specific single-cell dataset under investigation.

We also found that scRNA-seq and bulk RNA-seq data do not always correlate well for all gene pairs and explored different factors that could explain this. Part of the variable correlation could be explained by the sparsity of the single-cell data, as higher expressed gene pairs correlated better, but at least a few example cases showed the potential occurrence of Simpson’s paradox. With regards to cell type–composition, however, the FACS-sorted datasets did not correlate better with single-cell datasets than the whole-blood bulk dataset, which could either be caused by the smaller sample size of the single-cell data, technical changes introduced by FACS or specific differences in the (sub-)cell types, as we had naive CD4+T cells and classical monocytes (subsets of CD4+T cells and monocytes, respectively) for BLUEPRINT and ImmuNexUT that we tested for the single-cell data. Another interpretation is that scRNA-seq and bulk RNA-seq data capture different functional gene clusters, as a previous study showed in tumor samples (48). One possible explanation for this is that bulk and single-cell capture different sources of variability. Whereas single-cell data captures between-cell variability, bulk data captures between-person variability, which is affected by additional factors like genetics and environment. Therefore, a statistical framework combining both data types could be beneficial in the future.

Our study sheds light on several important considerations for future scRNA-seq study design regarding personalized network construction and co-eQTL mapping. Firstly, we showed that several factors, including cell number and gene selection, greatly influence the stability of co-expression patterns. We observed a clear trend indicating that a certain minimum number of cells from one individual is needed to achieve a stable co-expression pattern (**Figure 3d**). Secondly, we also explored factors influencing the number and quality of co-eQTLs. We showed that the number of significantly detected co-eQTLs can be greatly increased by either increasing the number of individuals or by increasing the number of cells per individual (**Figure 4c, d**). We believe that future larger single-cell datasets such as two very recent studies (49, 50) and the sc-eQTLGen consortium (51) will improve statistical power to identify more robust co-eQTLs.

Furthermore, we showed that a sophisticated filtering strategy of tested SNP–gene– gene triplets is essential to maximize the number of reliable co-eQTLs. However, we also suggest that the filtering strategy should be designed for the specific goals of the respective analysis. In this study, we systematically searched for robust co-eQTLs and adapted our strategy to balance the trade-off between achieving a stable co-expression pattern and enlarging the search space. For this reason, we first selected SNP–gene pairs and then used co-expression strength as an additional criterion rather than the very stringent expression cutoff criterion we used in our benchmarking analysis. In contrast, in our previous study (11), we focused specifically on co-eQTLs among the eQTLs that changed after pathogen stimulation and performed a strict pre-filtering for a highly targeted analysis. In the current study, we were, in particular, able to replicate the most significant co-eQTLs from the targeted analysis (**Supplementary Figure 30**). While the targeted analysis identified additional lower significance co-eQTLs that are below our much stricter multiple testing-corrected significance threshold, we were able to quantify the number of co-eQTLs more broadly for several additional SNPs and to include, for the first time, a comparison across cell types. In other cases, a selection of known TF–target pairs or pathway information could be desirable, e.g. for prioritizing TFs connected with diseases for experimental validation purposes.

We showed that the annotated co-eQTLs could identify potential direct regulators of the associated eQTLs as well as the affected biological processes, with several examples based on a combination of different enrichment analyses. We identified several TFs either directly as co-eGenes or via enriched binding sites among the co-eGenes of a SNP–eGene pair, providing potential regulatory mechanisms for explaining the co-eQTL. For the eQTL rs1131017–*RPS26*, six enriched TFs were themselves co-eGenes in CD4+ T cells, providing compelling evidence to support the hypotheses that direct regulators can be identified among co-eQTLs. Among these six TFs, five are associated with lymphocyte activation, further strengthening the connection of the eQTL with lymphocyte activation and through this to autoimmune diseases.

Another interesting aspect of the rs1131017–*RPS26* example is that we revealed a potential mechanism for a previously described GWAS signal by showing cell-type-specific genetic regulation of a multi-functional gene. By comparing T cells and monocytes, we identified that *RPS26* may be involved in two distinct biological functions. Interestingly, these two distinct functional co-eQTL clusters are characterized by opposite effect directions. Moreover, while *RPS26* showed enough variation to be picked up as an eQTL effect, it did not show high correlation with either gene cluster (**Supplementary Figure 31**), which may be why understanding its role in multiple functions has been challenging up to now (36, 52). We envision that more multi-functional eGenes could operate in such a cell-type-specific manner, with variation in expression that could be explained as the downstream consequences of many other conserved or highly co-expressed gene clusters, and this understanding could assist in interpreting GWAS signals. We also observed that different eGenes could have shared upstream genes/pathways as we identified four common immune-related co-eGenes associated with rs393727–*RNASET2* and rs1131017–*RPS26*, and both SNPs were in LD with immune diseases (T1D and CD), suggesting a shared upstream process for these two eQTL effects. By providing cell-type-specific gene regulation backgrounds through co-eQTLs, we expect more eQTLs and GWAS signals to be explained in the relevant cell type via future large-scale co-eQTL studies.

For the other co-eQTL examples, no enriched TF was in the co-eGene list, potentially because the TFs were not measured in the scRNA-seq datasets due to low expression. Here, the enrichment allowed us to still identify relevant TFs for further exploration. A third group of co-eQTL examples were supported by GWAS or GO enrichment analysis but not TF enrichment analysis. Here, the co-eGenes revealed part of the disease-relevant network, but we could not pinpoint the direct regulatory TFs. One explanation for this may be that our study is still underpowered to discover co-eGenes, while the enrichment strategy works best when there are a substantial number of co-eGenes as for rs1131017–*RPS26*. Based on our evaluation, we estimated that future studies with larger sample size and more cells will identify many more co-eQTLs (**Figure 4c,d**). This can help identify the direct regulators for some of our other examples, where the current enrichment analyses provided no clear interpretation, as well as co-eQTLs associated with other SNP–eGenes.

There are also several challenges to interpret the identified co-eQTLs. Firstly, as discussed earlier, it is difficult to determine the direct and indirect regulators that work through co-expression among correlated co-eGenes. This creates problems in using correlation-based metrics to quantify replication performance. For example, all the co-eQTLs we identified in B cells were associated with the rs1131017–RPS26 pair, making the correlation-based r_b_ measure invalid for this case. Also, to reduce the multiple testing burden, we only tested the top-SNP, a choice that could pose additional challenges for follow-up analysis such as colocalization to identify the causal SNP. Moreover, comparison of co-eQTLs between cell types remains challenging. We showed that the number of co-eQTLs is strongly driven by the number of cells (**Figure 4c**), so that it is not meaningful to only compare the absolute number of co-eQTLs between cell types in the current study. Furthermore, the sparsity of the single-cell data lead to the removal of many lowly expressed genes which, combined with the strict filtering our analysis required, meant only a small number of genes were tested in all cell types. In addition, sub-cell-type composition can introduce false positive co-eQTLs within a cell type if a genetic variant influences the sub-cell-type composition and one of the tested genes shows sub-cell-type-specific differences in expression. However, in our evaluation of classical and non-classical monocytes, we observed no strong confounding of monocyte co-eQTLs by the sub-cell types (**Supplementary Figure 24**). We also found several SNPs in LD with GWAS SNPs for traits such as monocyte counts, but there was no additional evidence that these SNPs have an effect on (sub-)cell type composition. Still, these analyses were limited to a smaller number of samples and a small number of cells in the sub-cell types, so we cannot exclude that some co-eQTLs were caused by sub-cell-type composition effects of the co-eQTL SNPs.

Several of the limitations of our current analysis will be overcome by on-going technological developments. First of all, we expect that follow-up analyses with larger sample sizes and more cells per person will identify many additional co-eQTLs. This can be further enhanced by improvements in single-cell technologies that lead to better capture efficiency of expressed genes. CITE-seq (53) and similar technologies (54, 55) allow improved cell type and sub-cell type classification that can show the effect of sub-cell type differences more accurately. The combination of multiple-omics, such as scRNA-seq, scATAC-seq and/or single-cell proteomics (56–58), will enable us to capture regulation happening outside the mRNA level, and lead to improved association analysis of gene pairs above standard Spearman correlation.

## Conclusion

Through our co-eQTL mapping strategy we identified a robust set of co-eQTLs that provides insight into cell-type-specific gene regulation and leads for future functional testing. Among these results, we uncovered a potential mechanism for a previously identified GWAS signal and a multi-functional gene. Our evaluation of different technical factors provides valuable suggestions for future experimental study design. We believe that more co-eQTLs will be uncovered by applying our general co-eQTL mapping pipeline to future large-scale scRNA-seq data. We envision that these co-eQTLs will in the future help to position eQTL and GWAS signals into cell-type-specific GRNs by annotating which regulatory edges are affected by which genetic variants. This knowledge is important for interpreting the effects of genetic variants in general, but also specifically for improve personalized medicine through better genetic risk prediction for diseases and personalized drug treatment based on genotype (51).

## Methods

### Single-cell datasets

Three different scRNA-seq datasets were included in this study, both for benchmarking the associations and for combined meta-analysis of co-expression QTLs. All five datasets from the three studies were generated from human PBMCs and are referred to by their first author: the Oelen dataset (n = 104 donors) (11), the van der Wijst dataset (n = 45 donors) (10) and the van Blokland dataset (n = 38 cardiac patients) (19). Further specifications can be found in **Supplementary Table 1** and the respective manuscripts. The processed versions of the datasets from the original publications were used, including quality control and cell type identification (for details, see the respective publications). The Oelen dataset also contains cells stimulated with different pathogens, but we only included the unstimulated cells in this analysis to improve comparison with the other datasets. For the van Blokland dataset, we included the data from the time point 6–8 weeks after the individual was admitted to the hospital for myocardial infarction, again to improve comparison across datasets.

For cell type classification, we took the annotation for the Oelen data from their original publication (11) and annotated the van Blokland and van der Wijst datasets using the Azimuth classification method (59). For Azimuth classification, we used the following settings: 1) the FindTransferAnchors function to find anchors using the reference from publication (59), normalization method “SCT”, reference reduction “spca” and first 50 dimensions and 2) the MapQuery function to annotate cell types using the same reference and parameters such as reference.reduction = “spca” and reduction.model = “wnn.umap”. We then compared the annotation from the Oelen publication and the Azimuth classification and found high correspondence (**Supplementary Figure 32**). For analyses using the sub-cell-type classification, we always refer to the Azimuth classification results.

### Single-cell co-expression

We calculated the Spearman correlation of gene pairs in the three different single-cell studies (Oelen dataset (11), van Blokland dataset (19) and van der Wijst dataset (10)) and then compared between datasets and 10XGenomics chemistry. In the benchmarking section, correlation was calculated separately per cell type but together over all individuals and only for gene pairs for which both genes were expressed in at least 50% of the cells from the respective cell type. For the comparison between two datasets, the gene pair–wise Spearman correlation values from each dataset were compared using Pearson correlation.

### Rho calculation

Rho proportionality was calculated using the “propr” function in R, from the “propr” package, with the symmetrized value set to true. We used the v3 unstimulated monocytes to compare the rho proportionality values to Spearman-rank correlations of the same data. We filtered out genes expressed in fewer than 5% of cells, leaving 8,634 genes to be assessed. Concordance between rho values and Spearman correlations was assessed with Pearson correlation.

We also explored rho proportionality values for very lowly expressed genes because the log-normalization of the method potentially introduces false associations for these genes (60). However, the computational demand to run the method was so high that we could not evaluate all expressed genes at once. Instead, we subsampled a set of 50 very lowly expressed genes (expressed in 0–5% of the cells) and 50 very highly expressed genes (expressed in at least 90% of the cells) and calculated the rho proportionality and Spearman correlation values for each combination of these 100 genes. We then compared gene pairs for which both genes were lowly expressed, pairs for which both genes were highly expressed, and mixed pairs, for which one gene was lowly and one highly expressed.

Alternative association metrics besides rho proportionality and Spearman correlation are discussed in **Supplementary Text**.

### Validation in bulk datasets

Spearman correlations from single-cell data were compared to Spearman correlations made with three different bulk datasets: the BLUEPRINT Epigenome consortium data (21), the ImmuNexUT dataset (22) and the BIOS dataset(2). For BLUEPRINT, we further removed the first principal component from the monocyte dataset to remove any uncorrected covariates. For the ImmuNexUT dataset, preprocessing was performed as described in the publication: we filtered out genes with less than 10 counts in 90% of the samples, performed TMM normalization with edgeR and scaling to CPM, batch corrected with combat and removed samples with a mean correlation coefficient smaller than 0.9. For the BIOS dataset, we corrected for 20 RNA Alignment metrics and then calculated the co-expression values using all individuals.

We then calculated the Pearson correlation across all gene pair-wise correlation values. As BLUEPRINT and ImmuNexUT are cell type-sorted datasets, we matched the cell types between bulk and single-cell data in these cases in the comparison. Again, we used only genes expressed in at least 50% of the cells from the cell type. This threshold was chosen after our initial evaluation of different thresholds from 10% to 90% in the comparison of BLUEPRINT and Oelen v3 dataset, with 50% chosen to balance the number of genes that can be used against the correlation strength between the datasets.

### Validation using CRISPR knockout data

To further validate the correlation values, we used CRISPR knockout data from (23). Mixscape was used to identify perturbed vs unperturbed cells for each CRISPR perturbation (27). We selected five knockout genes for which a sufficient number of successful CRISPR-perturbed cells were identified and that were expressed in our single-cell dataset (Oelen v3 dataset, CD4+ T cells) in > 50% of cells. The publication identified DE genes in wild-type vs perturbed cells and wild-type vs non-perturbed cells, as labeled by Mixscape. We selected a credible set of DE genes that were expressed in the single-cell dataset and significant in the wild-type vs perturbed cells but not in the wild-type vs non-perturbed cells. For this, we applied FDR-correction based on all genes expressed in the single-cell dataset. The correlation of these genes was compared to the correlation of non-DE genes, i.e. all other genes expressed in the single-cell dataset, using the Wilcoxon rank-sum test (one-sided test with “greater” in DE genes). The same test was done using the naive CD4+ T cells from the ImmuNexUT dataset.

### Validation using STRING annotations

Following the same approach used for the CRISPR knockout data, we explored if gene pairs whose proteins are interacting show higher correlation. We used the STRING database (version 11) (28), processed by the (18) benchmark study, to identify interacting gene pairs. We compared the correlation of gene pairs in STRING versus gene pairs not listed in STRING via Wilcoxon rank-sum test (one-sided test with “greater” for Gene pairs in STRING): once using the correlation estimates from the Oelen v3 dataset and once using the estimates from the ImmuNexUT dataset, both times for the CD4+ T cells and filtered for genes expressed in > 50% of single cells.

### Exploring Simpson’s paradox

To identify whether our strategy to identify single-cell co-expression is affected by Simpson’s paradox and whether bulk-based approaches would suffer from it, we studied the co-expression outcomes for two different strategies. In both strategies, we only included genes with non-zero expression in at least half of all monocytes in the Oelen v3 dataset. In the first strategy, we calculated Spearman correlations for gene pairs per individual separately for each gene pair. In the second strategy, we calculated the average expression of genes per individual and then calculated the Spearman correlation between genes. To identify potential Simpson’s paradox events, we looked into the gene pairs that had the largest deviation in co-expression estimate between the two strategies.

### Comparison between cell types

After successful validation of the Spearman correlation values, we compared differences between cell types within one dataset for the Oelen v2 and v3 dataset. Here we applied the same strategy as in the dataset comparison. We selected genes expressed in 50% of the cells from both cell types for each corresponding comparison, calculated Spearman correlation per gene pair within each cell type and followed up with Pearson correlation to compare both cell types. We also explored the absolute distribution of correlation coefficients between the cell types.

### Comparison between individuals

Again, we applied the same strategy as for the cell type and dataset comparison. We calculated gene pair–wise Spearman correlation values for each cell type and donor separately, taking all genes expressed in 50% of cells from the cell type in general (not per donor). We then compared each donor with each other donor by calculating Pearson correlation over the gene pair-wise correlation values to get a distribution of how well donors match per cell type.

To explore the effect of the number of cells per donor on this distribution, we subsampled each cell type to different numbers of cells (depending on the frequency of the cell type). For this, we take all individuals with at least this number of cells in this cell type and subsample the cell number to exactly this value for each individual. We stop subsampling at a threshold for the cell type when more than 75% of all measured individuals have fewer cells than the threshold. For the four most abundant cell types (CD4+ T cells, CD8+ T cells, monocytes and NK cells), we additionally fitted a logarithmic curve separately for each cell type to better quantify the connection: correlation_individuals ∼ log(number_cells) (with log being the natural logarithm). We then used the fitted formulas to extrapolate up to 1,500 cells for each cell type.

### Power calculation

For power calculation, we use an F-test, as implemented in (61), with a sample size of 173 (the total size of the combined cohorts), a heritability between 10% and 30% and a Bonferroni-corrected significance threshold of 0.05. The range for the heritability was chosen based on previously detected co-eQTLs (11). The number of tests influences the Bonferroni-corrected thresholds and depends on the selected gene–gene–SNP triplets. Here we assumed only one SNP per gene pair and all genes are tested against each other. Then, we increased the non-zero ratio threshold for gene selection from 0 to 0.95 (monocytes, Oelen v3 dataset), got the number of tests and calculated the power. Testing multiple SNPs per pair would further increase the total number of tests and reduce the overall power.

### eQTL mapping

We performed a meta-analysis to identify significant eQTL in four out of the five single-cell datasets (Oelen v2 and v3 dataset (11), van Blokland v2 dataset (19) and van der Wijst dataset (10)). We excluded the van Blokland v3 dataset because the sample size was so small that few variants lay above the MAF threshold (see below). Due to the limited sample size, we chose to perform a constrained eQTL mapping rather than a genome-wide mapping. To select the SNP–gene pair to test for eQTL mapping, we took the eQTL results from the largest meta eQTL analysis study in whole blood (2) and selected the most significant SNP for each gene. This resulted in 16,987 SNP–gene pairs to test. For these selected SNP–gene pairs, we performed eQTL mapping using eQTLPipeline v1.4.9 (62) within a *cis*-window of 100 kb, using 10 permutation rounds for determining FDR as described in (2) and a MAF of 0.1.

### Co-expression QTL (co-eQTL) mapping and the filtering strategy

First, we generated all possible combinations of the cell-type-specific eQTL findings (denoted as SNP–eGene) from the constrained eQTL mapping procedure in the respective cell type (as explained in the eQTL mapping method section above) and all other genes (denoted as co-eGene) that are expressed in the corresponding cell types. This resulted in the full list of SNP–eGene–co-eGene triplets for co-eQTL mapping analysis. We then calculated co-expression using Spearman correlation for the unique eGene–co-eGene pairs for each individual using untreated cells of the six major cell types (CD4+ T and CD8+ T cells, monocytes, B cells, NK cells and DCs) and the sub-cell-types in monocytes (classical monocytes and non-classical monocytes). For each gene pair, we counted the ratio of individuals who exhibit a significant correlation (nominal p-value from Spearman correlation < 0.05). If at least 10% of individuals showed a significant co-expression correlation for the specific eGene–co-eGene, we took this gene pair further into follow-up analysis. The total number of tests for each cell type can be found in **Supplementary Table 4**.

To assess the impact of cell numbers and sample numbers on the quality and quantity of co-eQTLs, we artificially created a few scenarios with fewer cells per individual and fewer individuals using a random subsampling strategy. To examine the impact of cell numbers, we randomly subsampled the CD4+ T cells per individual to three different levels (50, 150 and 250 cells). In each level, we kept the individuals with fewer cells, randomly subsampled those with a cell number higher than the corresponding level and performed the co-eQTL analysis using the strategies mentioned. Similarly, to examine the impact of sample numbers, we randomly subsampled 50 and 100 individuals, and excluded nine individuals with fewer than 10 CD4+ T cells for both scenarios.

### Multiple testing correction strategy for co-eQTL

To account for the correlation structure for gene pairs with one common gene and genome-wide, we modified and applied the permutation-based multiple testing correction strategy from fastQTL (29), implementing the method as follows. For each SNP–eGene–co-eGene triplet, we performed 100 permutations. Then, for each SNP– eGene pair, we determined the lowest p-values per permutation over all the genes (co-eGene) tested for the SNP–eGene pair. This resulted in the 100 lowest permuted p-values per SNP–eGene pair. For each SNP–eGene pair, we fitted a beta-distribution over the 100 permuted lowest p-values, which enabled us to subsequently establish the empirical p-value for the lowest non-permuted p-value. Through this procedure we ensured that under the null test statistic each SNP–eQTLGen pair has a uniform p-value distribution. Finally, for all SNP–eGene pairs, we calculated Benjamini-Hochberg FDR over the empirical p-values.

For each SNP–eGene pair, we also derived a p-value cutoff that indicates which of the co-eGenes are significant for that SNP–eGene pair via the following steps. After determining the FDR for all SNP–eGene pairs, we determined the empirical p-values that are closest to FDR = 0.05. Using the beta distributions for each SNP–eGene pair, we then determined its nominal p-value threshold. All co-eGenes with a nominal p-value lower than the corresponding p-value threshold for that SNP–eGene pair were considered significant.

### Replication in BIOS dataset

We replicated the co-eQTL findings in bulk whole-blood RNA-seq data from the BIOS Consortium, using the same method described in a previous study (8). Briefly, we implemented the following ordinary least squares model with the Python package statsmodels (63): *eGene ∼ SNP + co-eGene + SNP:co-eGene*. We then examined the effect sizes of the interaction term SNP:co-eGene and used Benjamini-Hochberg procedures for multiple testing correction.

### Calculation of r_b_ values and allelic concordance

We used the same evaluation metrics to quantify the cell-type-specificity and replication performance in the BIOS data set of the co-eQTLs. First we used the rb method with modification. We followed the same procedures as the original study (31) but chose a suggested alternate strategy to estimate errors across gene pairs between two tissues. Whereas the original paper used null SNPs per each eQTL for this purpose, we tested only the significant eQTL SNP for SNP–eGene–co-eGene triplets and therefore we did not have information for the null SNPs. Thus, we used the alternative approach indicated in the original paper with **Equation 1)**, where *r_e_* is the estimation errors across gene pairs between two tissues, *r_p_* is the correlation of co-expression levels between two cell types in the overlapping sample, *n_s_* is the number of overlapping samples, *n_i_* and *n_j_* are the number of samples in cell typed *i* and *j*, respectively. For the BIOS replication, we excluded overlapping individuals from the BIOS RNA-seq dataset for the replication analysis. Additionally, in cases where fewer than 10 co-eQTLs were tested in the replication analysis, we could not get a robust estimation of the r_b_ value and hence represent them as NAs in the results section.

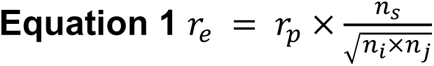

Due to our filtering strategy, we did not always test the same set of SNP–eGene–co-eGene triplets in all cell types. Therefore we also need to compare the tested ratio when quantifying the cell-type-specificity. The tested ratio means the SNP–eGene–co-eGene triplets that were also tested in another cell type or the BIOS replication analysis. A third evaluation metric that we used is allelic concordance between the discovered co-eQTLs and the results in the replication study. This is defined as the ratio of co-eQTLs with concordant effect direction by the number of significant co-eQTLs identified in the replication study.

### Biological interpretation based on enrichment of GO terms, TF binding sites and GWAS variants

We explored the biological function of the co-eQTLs based on different enrichment analyses that all tested if co-eGenes associated with the same SNP–eGene pair in the same cell type show similar functional properties. For this, we selected all SNP–eGene pairs that had at least five significant co-eGenes in the same cell type.

First, we performed GO enrichment analysis separately for each co-eGene set, grouped by SNP–eGene and cell type, applying the R package clusterProfiler (ver 4.0.5) (64) and performing FDR multiple testing correction separately for each SNP–eGene pair across the different GO terms (defining enrichment below FDR < 0.05 as significant). As the background set for the enrichment, we used all genes tested in the co-eQTL analysis in the respective cell type.

Next, we explored if these co-eGene sets were enriched for certain TF binding sites. TF annotations were taken from ChIP-seq peaks processed in the ReMap 2022 database (33), which we filtered for cell lines associated with blood cell lines. We tested the overlap of these peaks with the promoter regions of the co-eGenes tested, defining the promoter region as the region 2kB upstream and downstream of the first transcription start site of the gene. Enrichment was tested based on Fisher’s exact tests for each TF, using all genes tested in the co-eQTL analysis in the respective cell type as the background set. We performed FDR multiple testing correction separately for each SNP–eGene pair over all TFs (defining enrichment below FDR < 0.05 as significant). Furthermore, we explored if the enriched TF itself was a co-eGene associated with the respective SNP–eGene pair and if the co-eQTL SNP or a SNP in high LD (R^2^≥ 0.9) lies in a binding site of the enriched TF. The SNPs in high LD were obtained from SNiPA (65) using the variant set from the 1000 Genomes Project, Phase 3 v5, European population, Genome assembly GRCH37 and genome annotations from Ensembl 87. For the GWAS annotations, we considered two different strategies. In the first approach, we annotated SNPs or SNPs in high LD (R^2^ ≥ 0.8) with GWAS loci from the GWAS Catalog (1), with the last updated timestamp being 3/1/2022, 07:13 AM (GMT+0100). LD information for this was taken from LDtrait (66) with the following parameters: window size = 500KB, reference population = 1000 Genomes CEU, GRCh37). In the second approach, we used the magma method (67) to assess enrichment of GWAS associations among co-eGenes. We obtained uniformly processed GWAS summary statistics for 114 traits that were used for the GWAS analysis of the GTEx consortium (68, 69). We then followed the strategy previously described by (67). We defined gene sets for each co-eQTL SNP in each tissue as the set of significant co-eGenes associated with the SNP, as done for the GO and TF enrichment analysis. Protein names/gene symbols were converted to Entrez gene ids and mapped to the corresponding annotations on the human genome assembly 38. We performed individual magma analyses for each trait based on summary statistics and LD structure from the 1000 genomes European reference panel for all gene sets compared to the background set of genes tested for co-eQTL, always conditioning on default gene-level covariates (for example, gene length). Subsequently, we applied the Benjamini-Hochberg method and selected gene set–trait associations with FDR < 5%.

After we observed different distributions of co-eQTLs for rs11311017–*RPS26* with regards to the direction of effect in the different cell types, we repeated all enrichment analysis (GO, TF and GWAS) separately for the positively associated co-eGenes and negatively associated co-eGenes in CD4+ T cells.

### Direction of effect

We compared the direction of effect in eQTLs and co-eQTLs by comparing the direction of the zscores. After ensuring that the reference allele aligns in the eQTL and co-eQTL analysis, co-eQTLs for which the sign of the zscore matches the sign of the eQTL zscore are called concordant. If otherwise, they are called discordant.

## Declarations

### Ethics approval and consent to participate

Ethics approval was requested and approved for each of the data sets. The Lifelines DEEP study was approved by the ethics committee of the University Medical Centre Groningen, document number METC UMCG LLDEEP: M12.113965. All participants signed an informed consent prior to study enrollment. All procedures performed in studies involving human participants were in accordance with the ethical standards of the institutional and/or national research committee and with the 1964 Helsinki declaration and its later amendments or comparable ethical standards.

### Consent for publication

All authors have read and approved the submission of this manuscript.

### Availability of data and materials

All code is available on github at https://github.com/sc-eQTLgen-consortium/co-expressionQTLs

BIOS: https://www.bbmri.nl/acquisition-use-analyze/bios

Oelen datasets: https://eqtlgen.org/sc/datasets/1m-scbloodnl.html

Van Blokland data set: manuscript in preparation.

Van der wijst data set: https://eqtlgen.org/sc/datasets/vanderwijst2018.html

### Funding

Horizon2020 №860895 (MK)

NWO-VENI 192.029 (MW)

NWO-VICI, 917.14.374 Oncode Investigator grant (LF)

NWO-VIDI grant number 917.164.455 (MAS)

Chan Zuckerberg Initiative grant number 2019-202666 (MH)

### Authors’ contributions

SL, KTS, DV, and MK implemented the pipeline, analyzed the data, and drafted the manuscript. HW implemented the QTL mapping and permutation tool. LF, MH, MW, HW, and MAS contributed to the project design, results interpretation, and supervision of the work. RO, IB, HG, PH, and MW provided the data. IB, HG, and PH generated the van Blokland data set. All authors reviewed the manuscript and approved the final manuscript.

## Acknowledgements

We are very grateful to all the volunteers who participated in this study. We thank Kate Mc Intyre for editing the manuscript systematically and extensively.

We thank Martijn Vochteloo for the BIOS replication data preparation.

We thank the UMCG Genomics Coordination Center, the UMCG Research IT programme, the UG Center for Information Technology and their sponsors BBMRI-NL & TarGet for storage and compute infrastructure. We thank the Biobank-Based Integrative Omics Studies (BIOS) Consortium, funded by the Biobanking and Biomolecular Research Infrastructure Netherlands (BBMRI-NL), a research infrastructure financed by the Netherlands Organization for Scientific Research (NWO) under award number 184.021.007.

