## Supplementary Text for "Identification of genetic variants that impact gene co-expression relationships using large-scale single-cell data"

### MetaCell

As we observed a decline in concordance with bulk data sets for genes with more zero values (**Figure 2c**), we looked into strategies to limit the sparseness of the data. One possibility would be to impute missing values. However, as this tends to lead to false positive associations, we decided against it (1,2). As an alternative, we merged similar cells into non-overlapping *MetaCells* (3) to reduce the zero-genes at the cell level.

#### Methods

We tested the *MetaCell* algorithm on the Oelen v3 dataset, Monocytes, running the algorithm separately per sample and calculating the average expression per gene over all cells assigned to the same metacell. We ran the algorithm on the full dataset, including also the stimulated conditions to increase the heterogeneity among the cells and so improving the clustering. As expected, meta-cells could be always clearly assigned to one condition, as cells from the same condition clustered together. For the evaluation shown in **Supplementary Figure 3**, only the untreated meta-cells were used.

Using the same approach as before, we evaluated the Spearman correlation from the meta-cells with the Spearman correlation from the BLUEPRINT dataset (4) and compared the outcome with the concordance when using the single-cell data directly. We tested different parameters to reduce the granularity of the meta-cells by changing the parameters the  K and min_mc_size in the function mcell_mc_from_coclust_balanced (K between 5 and 20, min_mc_size between 3 and

10), which had however nearly no impact. As the parameter had only a small influence in the granularity of the metacells, we additionally tested meta-cells generated from Leiden clustering with a resolution of 20 (5) and performed the same evaluation.

#### Results

The original *MetaCell* algorithm provided too few meta-cells for a proper calculation of correlation between genes per cell type, so we implemented our own adaptation based on Leiden clustering (see Methods). We saw clearly that the fraction of zero values in the meta cells declined. However, this led to far fewer data points (metacells) per individual and could not in the end increase the concordance with bulk data from BLUEPRINT (**Supplementary Figure 3**). For this reason, we did not proceed with the *MetaCell* idea, although a more extensive exploration of this approach in future work could still lead to a promising alternative strategy.

### Alternative GRN construction methods

Apart from Spearman correlation, we also tested other gene regulatory construction methods, namly Rho proportionality measure (6), and GRNBoost2 (7). Several other methods developed for single cell data, could not be tested, as we could not infer a reliable pseudotemporal ordering from our data set, for which we tested RNA velocity (8) and SCORPIUS (9). We discussed the results from Rho proportionality in the main manuscript, the other approaches are explained here.

#### Methods

**GRNBoost2**

We took the genes that were expressed in at least 50% of the unstimulated monocytes in the Oelen v2 data, and calculated the edge weight between the gene pairs with GRNBoost2. We repeated this analysis using monocyte data from Blueprint. Then we correlated the edge weight for all gene pairs from Oelen v2 data and that from Blueprint. We implemented the GRNBoost2 analysis using the tool and container provided in the benchmark study (10).

**RNA velocity**

For the RNA velocity estimate, we used *velocyto* (11) to get both spliced and unspliced gene count matrices followed by *scVelo* (8) for the velocity. scVelo was run on the combined set of untreated and stimulated cells, filtered for the subset of classical monocytes, using the dynamical mode and the 2000 highest variable genes.

**SCORPIUS**

For this analysis we selected the classical monocyte (12) to ensure that sub cell type composition will not confound the final results. We selected cells from 3 conditions: untreated condition, stimulated condition with Candida for 3 hours and stimulated condition with Candida for 24 hours. We inferred the pseudotime ordering of the cells with the scripts provided along with the study (9) with default settings.

### Results

Additionally to Rho proportionality and Spearman correlation(see main manuscript), we tested other GRN construction methods suggested in a recent benchmark paper (10). For algorithms that do not require time-stamps for cells, we tested the scalable top-performing method suggested in the benchmark study (10) named GRNBoost2 (7). However, we observed poor correlation (spearman r = 0.17) between the GRN inferred from BLUEPRINT data and that from our single cell data (**Supplementary Figure 5**) using GRNBoost2, which is much lower than that for Spearman correlation. Therefore, for the rest of this study we continued our analysis with Spearman correlation.

Several single-cell-specific GRN reconstruction methods had to be excluded, because they required pseudotemporal ordering of cells, which could not be reliably inferred from our data set. RNA velocity generally does not work well in blood data sets (8). To better explore the inferred dynamics, we included the pathogen-stimulated timepoints from the Oelen data set in our analysis (3h and 24h after pathogen stimulation), but the drastic expression changes after pathogen stimulation led to completely separate states instead of temporal trajectory (**Supplementary Figure 4**). For our datasets, the inferred pseudotemporal ordering of cells is algorithm-dependent. We compared the ordering from RNA velocity (8) and a pseudotime ordering algorithm called SCORPIUS (9). Timestamps predicted by these two algorithms are poorly correlated (**Supplementary Figure 4b,c**) with each other. However, the ordering from RNA velocity, which should excel in inferring cell trajectories compared to algorithms solely based on transcriptomic similarity, does not correlate well with our experimental sampling time points (**Supplementary Figure 4a,b**), which makes the cell ordering results difficult to interpret.

### Additional promising co-eQTL examples

Another set of co-eQTLs partly supported by our enrichment analysis is the co-eQTLs associated with rs4147638 - *SMDT1* identified in CD4+ T cells, which were found to be enriched as for several GO terms including translation initiation and protein targeting to endoplasmic reticulum (**Supplementary Table 14**) and target for 5 TFs in CD4+ T cells and 37 TFs in CD8+ T cells (**Supplementary Table 15**). Similarly, in DCs, we identified 30 co-eGenes for the type 2 diabetes (T2D) SNP rs7935082 and eGene *MS4A7*, and these co-eGenes were enriched for T2D (**Supplementary Table 17**) and several endoplasmic reticulum associated GO terms (**Supplementary Table 14**).
