## Supplementary Figures and Supplementary Tables for "Identification of genetic variants that impact gene co-expression relationships using large-scale single-cell data"

### Supplementary Information

#### Supplementary Tables

***Supplementary Table 1.*** *Data set specifications*

***Supplementary Table 2.*** *Summary statistics of the eQTLs for the major cell types and meta-analysis*

***Supplementary Table 3.*** *eQTLs for major cell types*

***Supplementary Table 4.*** *Summary statistics of the co-eQTLs for the major and sub cell types without the filtering strategy (including the sub-sampling strategies and sub cell types)*

***Supplementary Table 5.*** *Summary statistics of the co-eQTLs for the major cell types with the filtering strategy (including the sub-sampling strategies and sub cell types)*

***Supplementary Table 6.*** *The significant co-eQTLs for the major cell types with the filtering strategy and their replication results in other cell types*

***Supplementary Table 7.*** *Summary of replication rb values for filtered co-eQTLs in other cell types*

***Supplementary Table 8.*** *Summary of replication rb values for filtered co-eQTLs in BIOS data*

***Supplementary Table 9.*** *BIOS replication for filtered co-eQTLs for major cell types*

***Supplementary Table 10.*** *The significant co-eQTLs for the major cell types without the filtering strategy and their replication results in other cell types*

***Supplementary Table 11.*** *Summary of replication rb values for unfiltered co-eQTLs in other cell types*

***Su[plementary Table 12.*** *BIOS replication for unfiltered co-eQTLs for major cell types*

***Supplementary Table 13.*** *Summary of replication rb values for unfiltered co-eQTLs in BIOS data*

***Supplementary Table 14.*** *GO enrichment for co-eGenes associated with the same eQTL, for all eQTLs with at least 5 co-eGenes*

***Supplementary Table 15.*** *TF enrichment of co-eGenes co-eGenes associated with the same eQTL, for all eQTLs with at least 5 co-eGenes, using the Remap 2022 ChIP-seq data*

***Supplementary Table 16.*** *Annotated co-eQTL SNP with GWAS results*

***Supplementary Table 17.*** *Enrichment of co-eGenes for GWAS results*

### Supplementary Figures

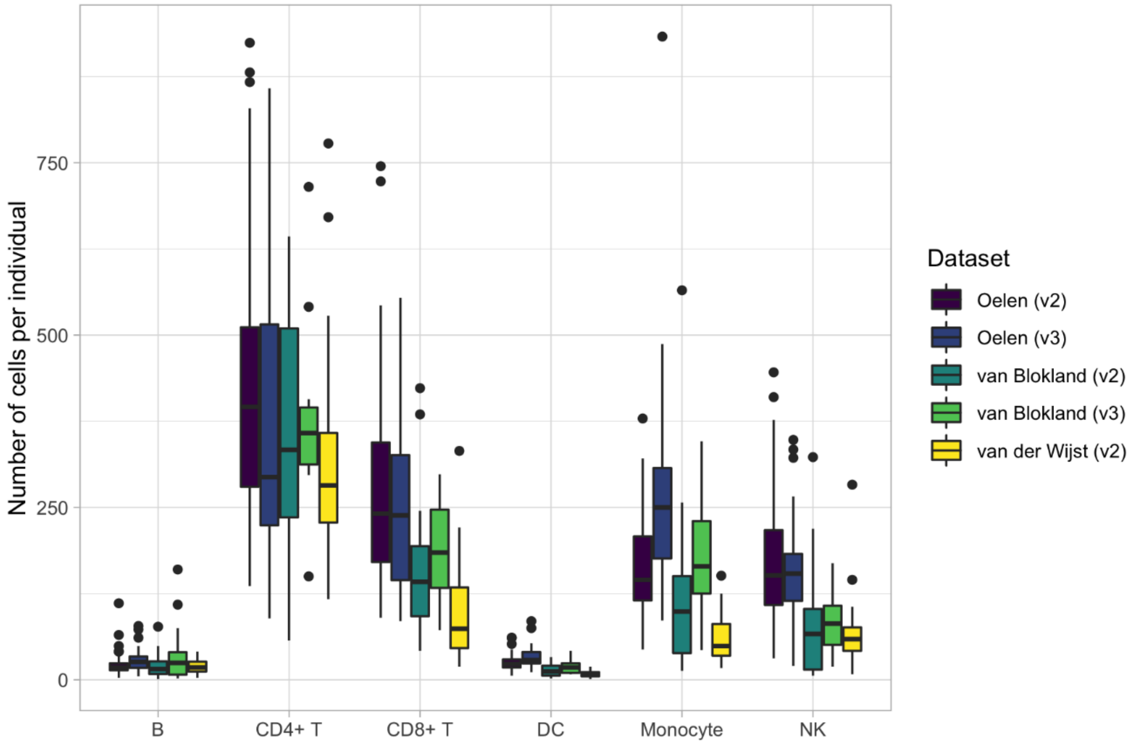

**Supplementary Figure 1.** *Number of cells per cell type and individual for each dataset*

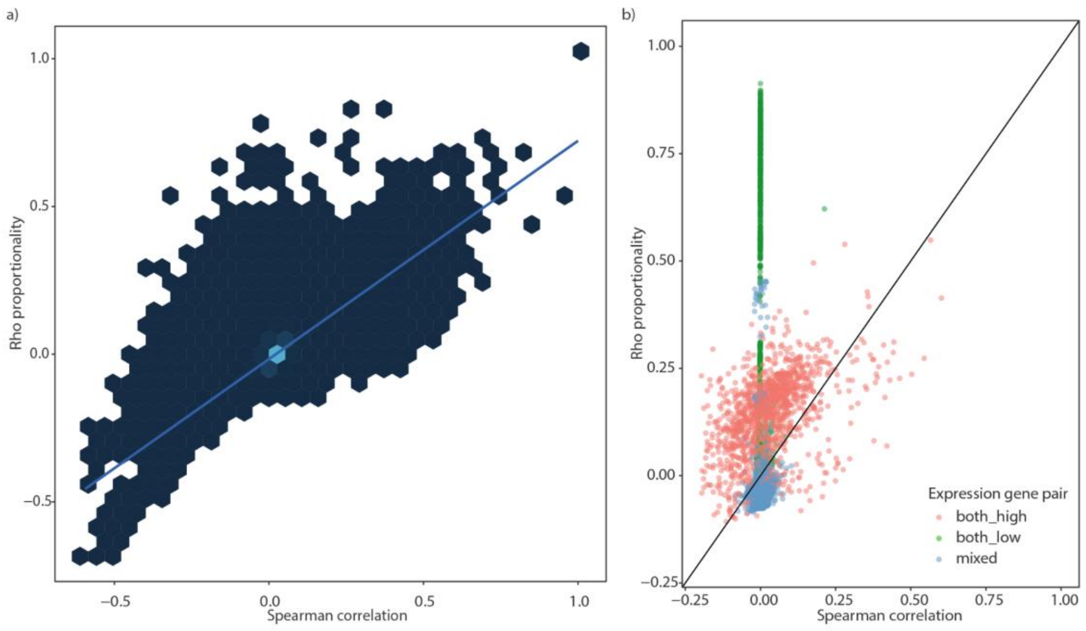


**Supplementary Figure 2.** *Comparison between proportionality (Rho) and Spearman correlation for genes that are expressed in at least 5% of the monocytes in untreated status. The color indicates the density. Light color corresponds to higher density.*

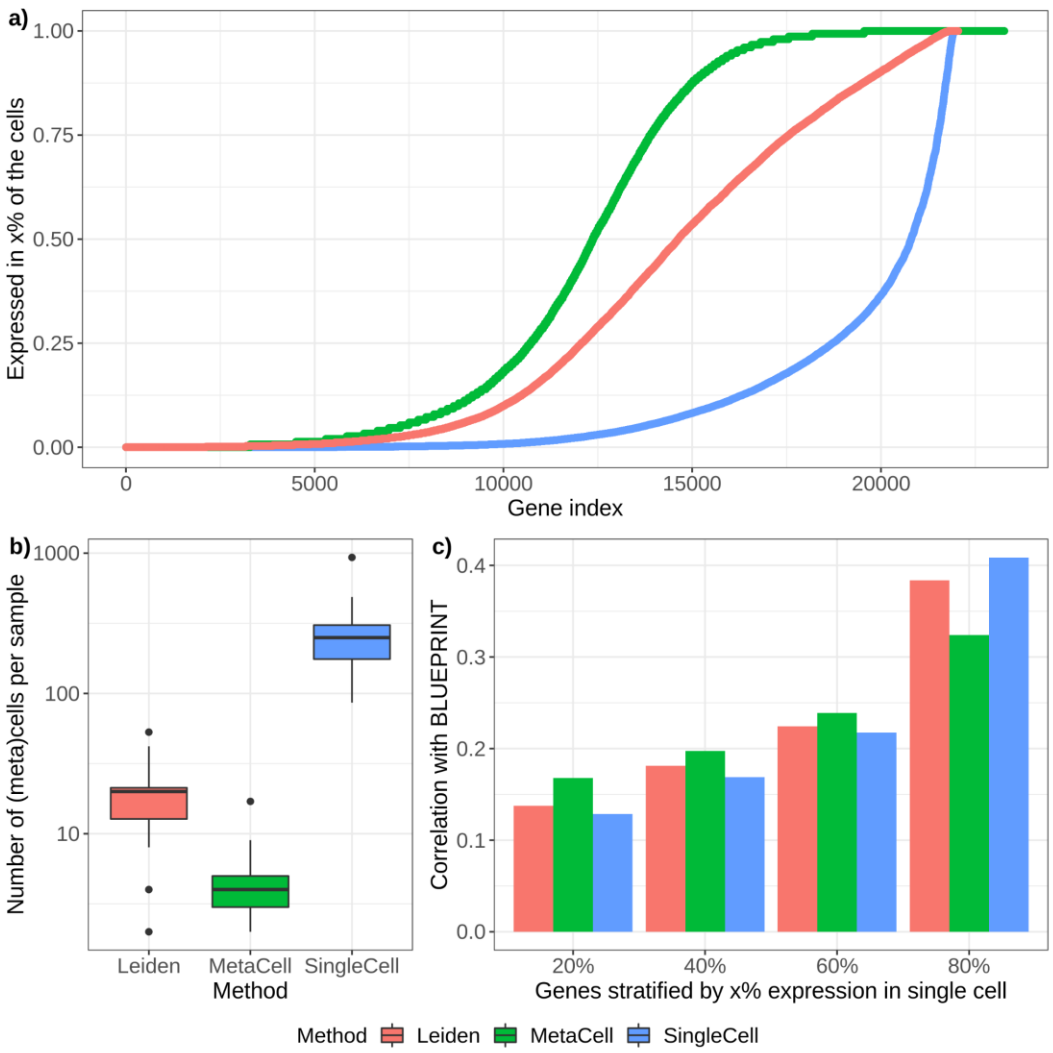


***Supplementary Figure 3.*** *Grouping cells to meta-cells*

*Similar cells were grouped to meta-cells, using either the original MetaCell algorithm (parameters shown in the plot: K=20, minCells=10) or our own implementation based on the Leiden algorithm (parameters shown in the plot: resolution=20) (see Methods for detail). All methods were applied to the Oelen v3 dataset, Monocytes.* ***a)*** *Both meta-cells generated from Leiden clustering and from the MetaCell algorithm lead to more genes expressed in at least x% of the cells compared to the original single cell data (visualized here via a cumulative density function).* ***b)*** *In contrast, the number of (meta)cells per sample is reduced with both algorithms drastically, this way reducing the number of measurement points to infer the correlation per sample.* ***c)*** *To* benchmark the performance, the correlation with the BLUEPRINT bulk data set was calculated (compared with Main Figure 2b). To evaluate how lower expressed genes are affected by the meta-cell grouping, the correlation is calculated separately for gene pairs were the non-zero expression level of both genes is between 20%-40%, 40%-60%, 60%-80% and 80%-100% of the cells (showing the first number on the x-axis).*

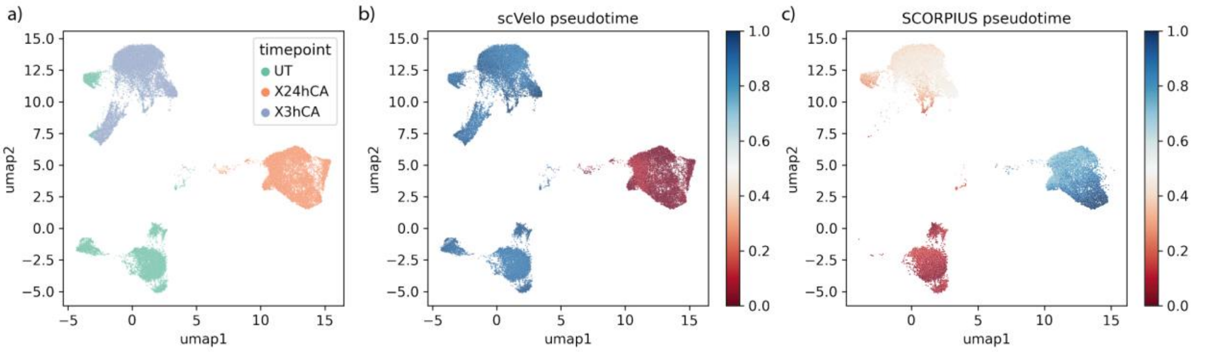


***Supplementary Figure 4.** Comparison for the inferred time-stamp information from RNA velocity with scVelo and pseudo-time ordering algorithm SCORPIUS, and the sample time for the single cell data collection during stimulation experiments*

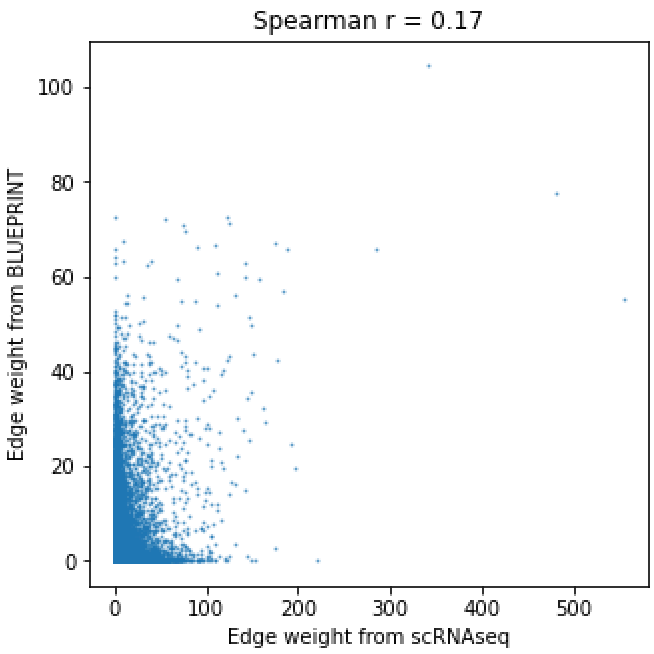

***Supplementary Figure 5.*** *Prediction performance comparison for STRING gene pairs between single cell monocytes and BluePrint monocytes, for both data, the predicted gene pairs are from GRNBoost2 method*

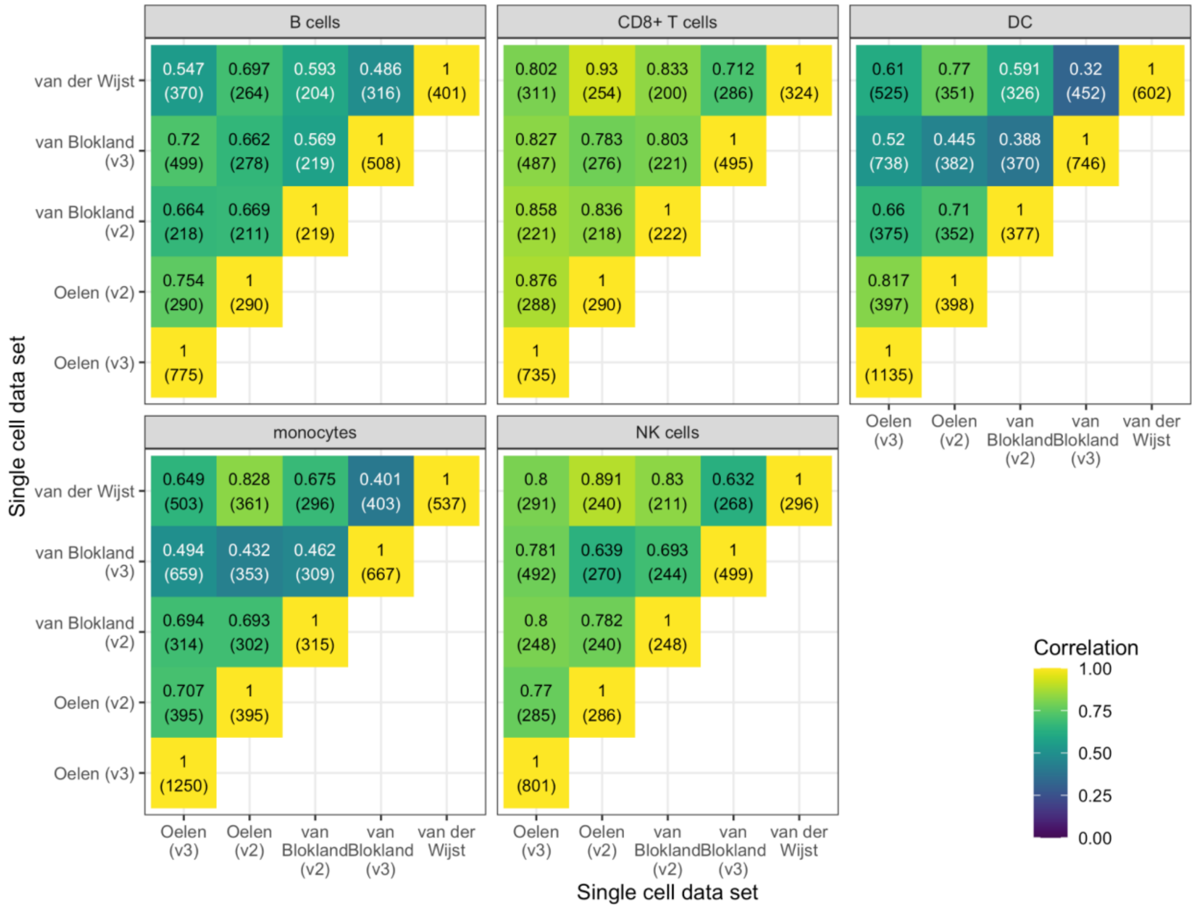


***Supplementary Figure 6.*** *Extension of Main Figure 2a for other cell types, comparing different single-cell datasets. Spearman correlation of the the Oelen v3 and v2 datasets, the van Blokland v2 and v3 datasets and the van der Wijst dataset were compared with each other, taking genes expressed in at least 50% of the cells in the corresponding datasets.*

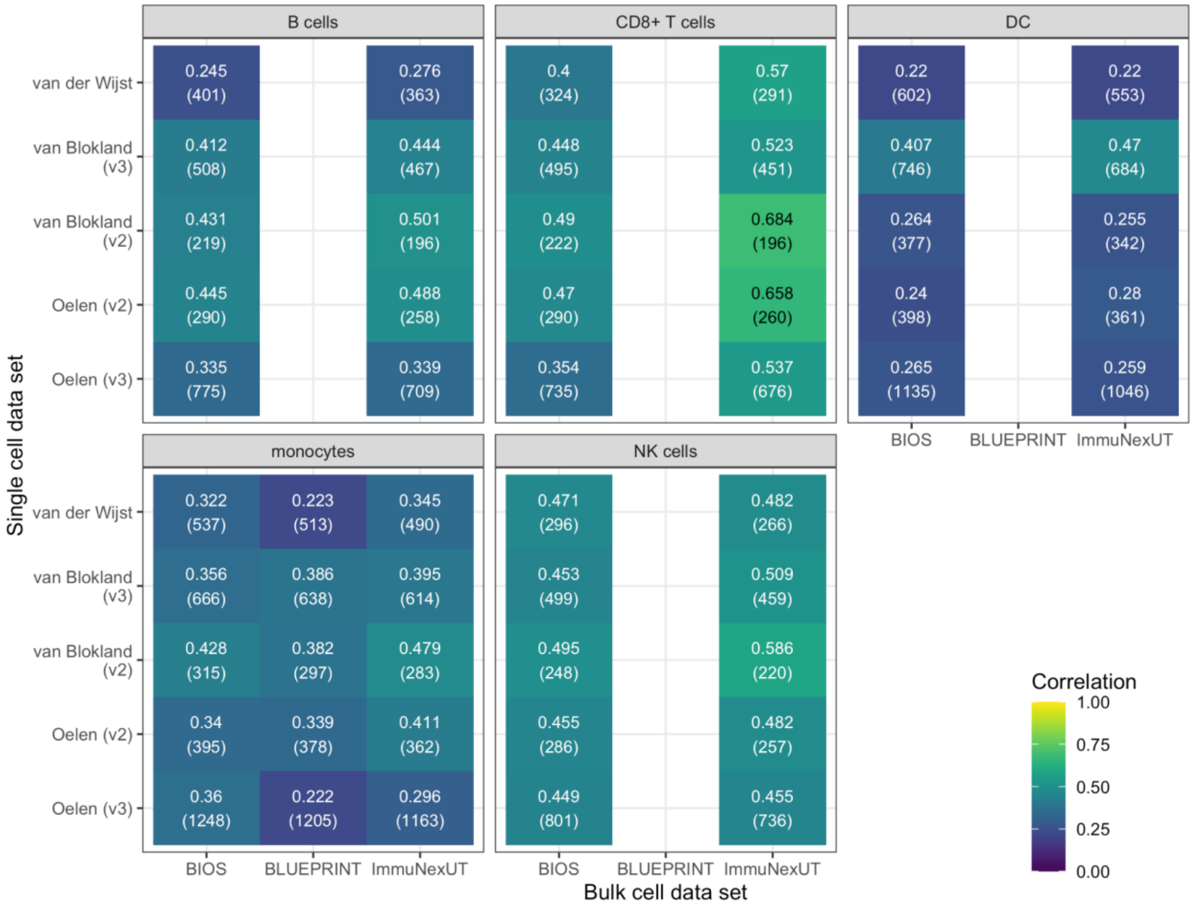


***Supplementary Figure 7.*** *Extension of Main Figure 2b for other cell types: Comparison of the co-expression profiles between the single-cell datasets with the bulk RNA-seq datasets from BLUEPRINT, ImmuNexUT (both measuring FACS sorted cell types) and BIOS (whole blood). For BLUEPRINT, classical monocytes were measured, for ImmuNexUT, the compared cell types were naive B cells, naive CD8+ T cells, myeloid DCs, classical monocytes and NK cells. Again only genes were taken that were expressed in at least 50% of the cells for the single-cell dataset. The number of tested genes is shown in brackets in each square below the exact Spearman correlation value.*

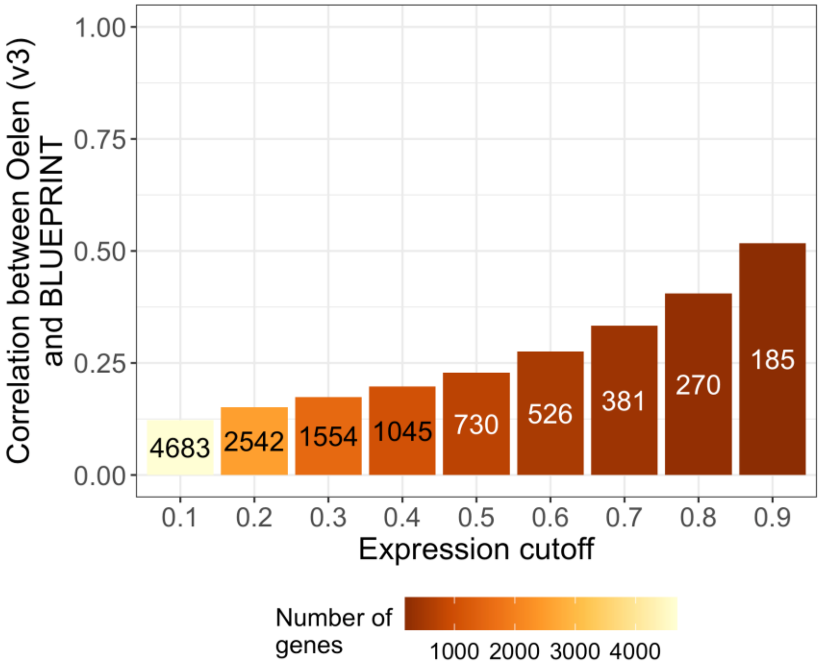


***Supplementary Figure 8.*** *Analysis from Main Figure 2c with the BLUEPRINT dataset instead of the ImmuNexUT dataset: Relationship between the co-expression similarity between the BLUEPRINT naive CD4+ T cells and Oelen v3 dataset CD4+ T cells and increasing gene expression cutoffs (the ratio of cells with non-zero expression for a given gene). Both the color scale and the numbers in the bar plot show the number of tested genes.*

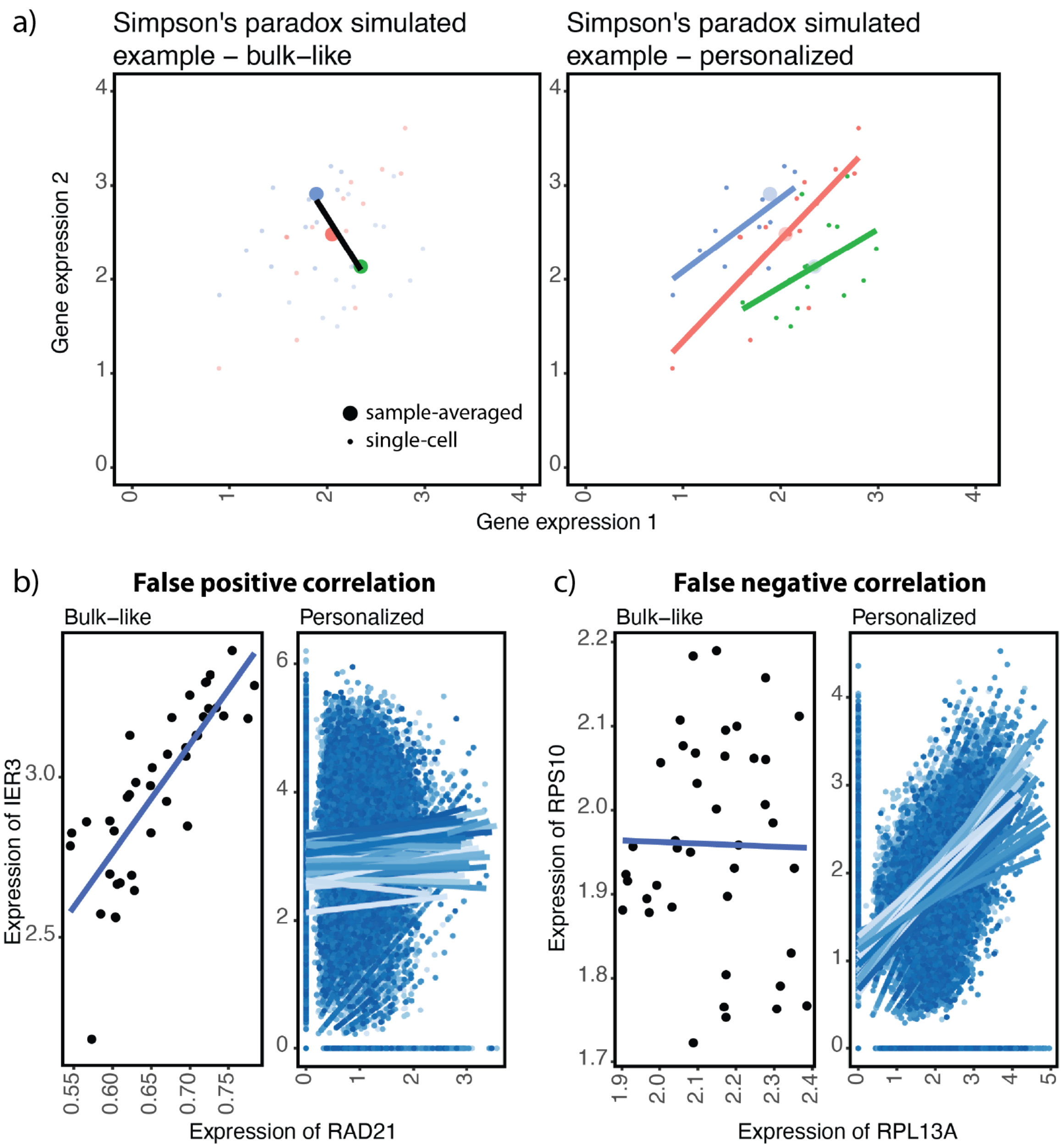


***Supplementary Figure 9. Simpson's paradox in expression data a)*** *A simulated example showing how Simpson's paradox can appear in expression data. The colors of the dots represent the sample. Each small dot represents a cell and the large dots represent the average expression across cells for that individual. The line in the left figure is the regression line for the sample-averaged dots and the lines on the right are the regression lines for all cells for each of the three samples.* ***b)*** *An example showing a false positive correlation identified by the bulk-like expression data but not identified in the personalized manner* ***c)*** *An example showing a false negative correlation not*

*identified if aggregating the scRNA-seq data with the bulk-like approach, but a true correlation identified by the personalized expression data.*

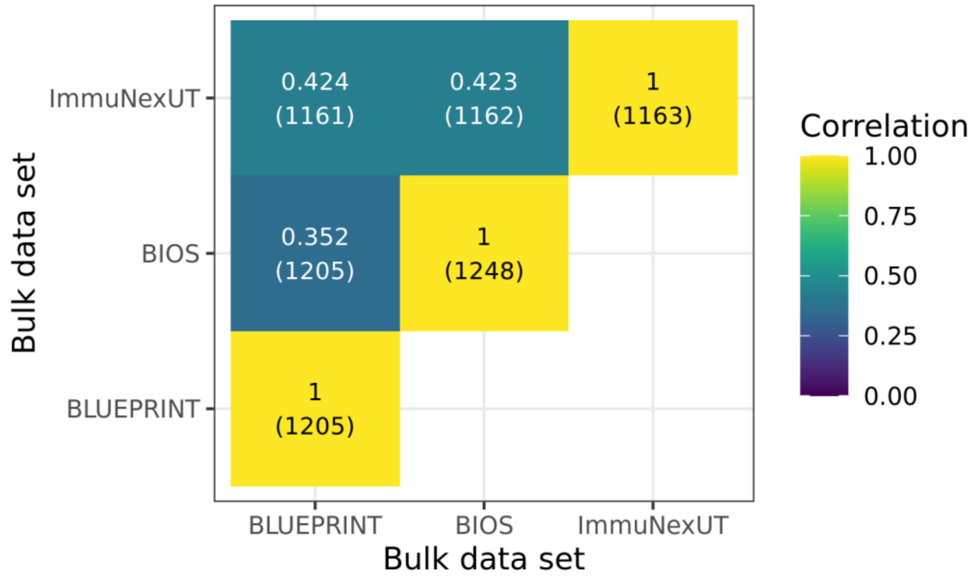


***Supplementary Figure 10.*** *Extension of Main Figure 2d for monocytes: Comparison of the co-expression profiles between the bulk RNA-seq datasets from BLUEPRINT, ImmuNexUT (both measuring FACS sorted classical monocytes) and BIOS (whole blood). In all datasets, only genes expressed in 50% of the cells from the Oelen v3 dataset were selected, to make it comparable with Supplementary Figures 6 and 7. The number of tested genes is shown in brackets in each square below the exact Spearman correlation value.*

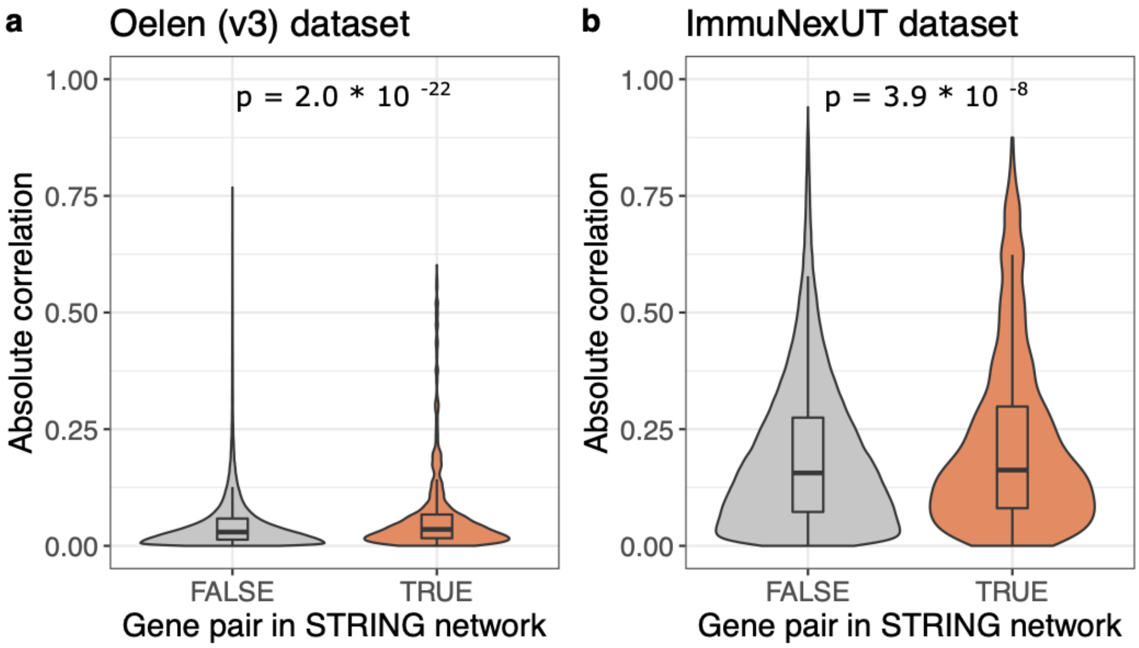


***Supplementary Figure 11**. Enrichment of correlated genes among gene pairs whose proteins are interacting according to the STRING database, taking correlation values from Oelen v3 single-cell dataset in **a)** and ImmuNexUT bulk dataset in **b)**. P-values in the plot show the significance level of the Wilcoxon rank sum test.*

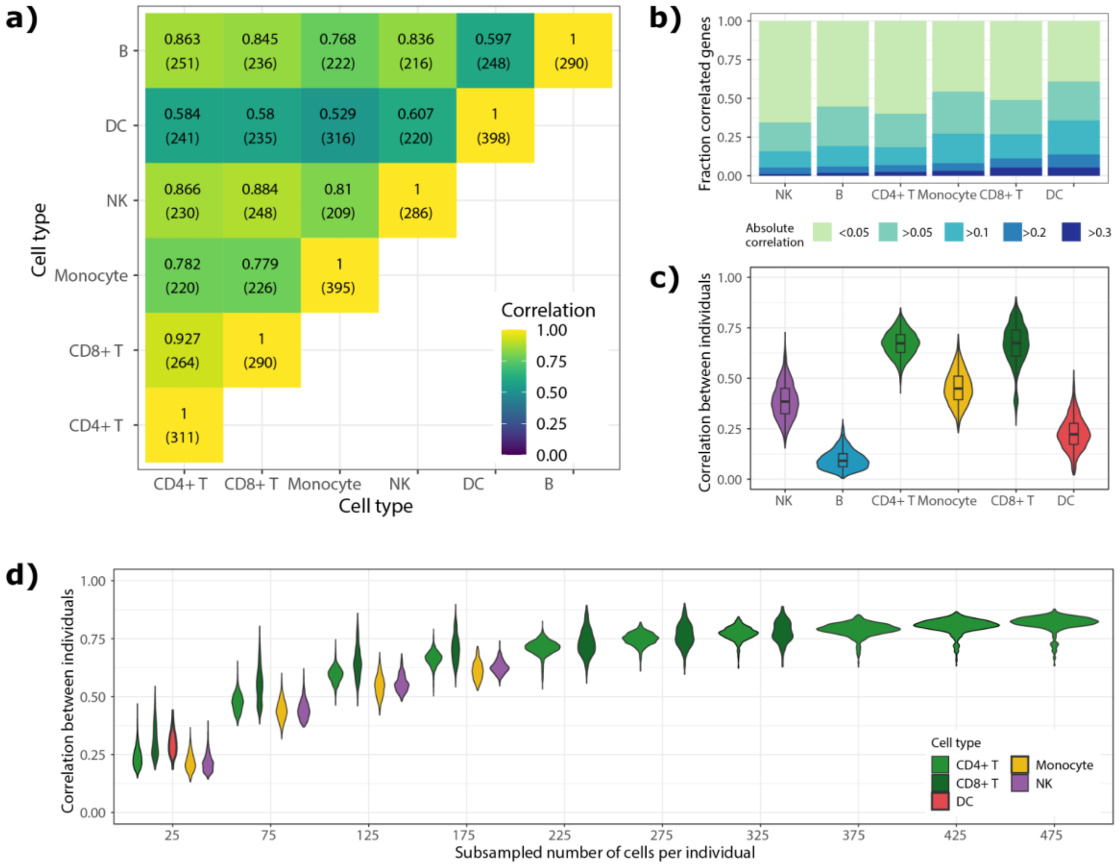


***Supplementary Figure 12.*** *Adaptation of Figure 3 for Oelen v2 dataset instead of Oelen v3 dataset, showing the same trends: Each analysis was done for all gene expressed in at least 50% of the cells for the respective cell type.* ***a)*** *Comparing co-expression patterns across cell types within the Oelen v2 dataset, for genes expressed in 50% of the cells for both cell types in each pairwise comparison (same approach as in Figure 2a-c)* ***b)*** *Correlation distribution within each cell type* ***c)*** *Correlation between different individuals within each cell type, showing the distribution of all pairwise comparisons between individuals.* ***d)*** *Dependence of number of cells on the correlation between individuals, separately for each cell type. In each subsampling step, all individuals are taken that have at least this number of cells and subsampled to exactly the number (this leads to removal of some individuals for higher number of cells). B cells were not frequent enough to evaluate it in this data set.*

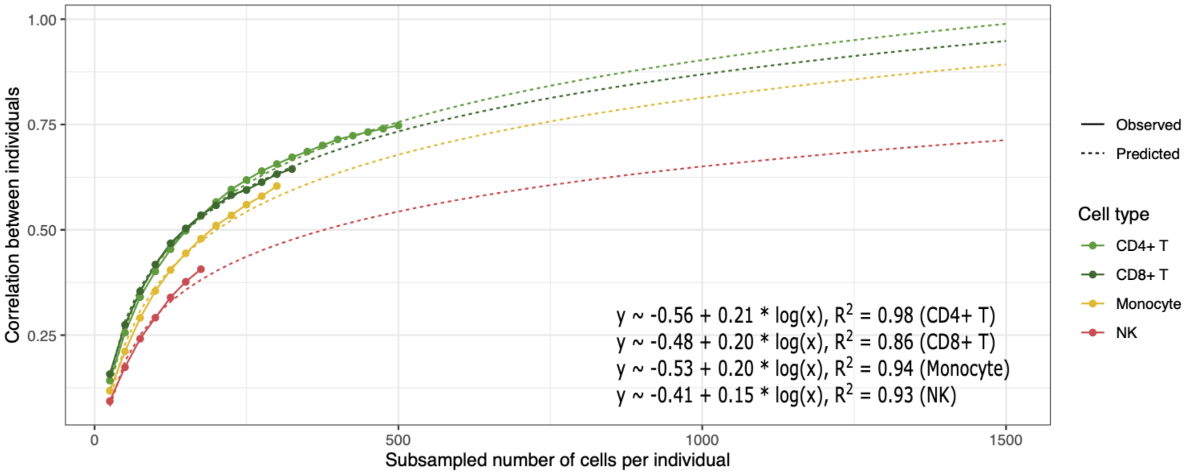


***Supplementary Figure 13.*** *Fitting a logarithmic curve (based on the natural logarithm) for the four most frequent cell types (CD4+ T cells, CD8+ T cells, monocytes, NK cells) to explain the correlation value between individuals by the number of cells per individuals (estimated curves and adjusted $R^2$ values for each cell type in the text). Dotted line shows extrapolation of this fit to predict correlation when increasing the number of cells up to 1,500 cells per individual and cell type.*

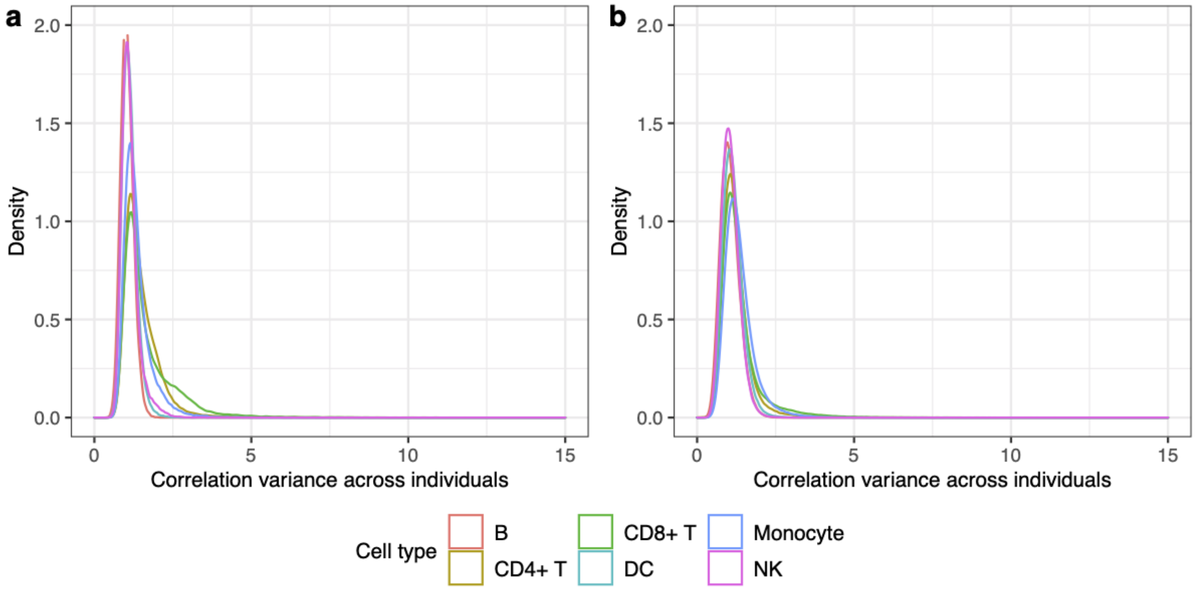


***Supplementary Figure 14.*** *Variance of gene pairs (correlation z-scores) across individuals per cell type for Oelen v2 dataset in* ***a)*** *and Oelen v3 dataset in* ***b)****.*

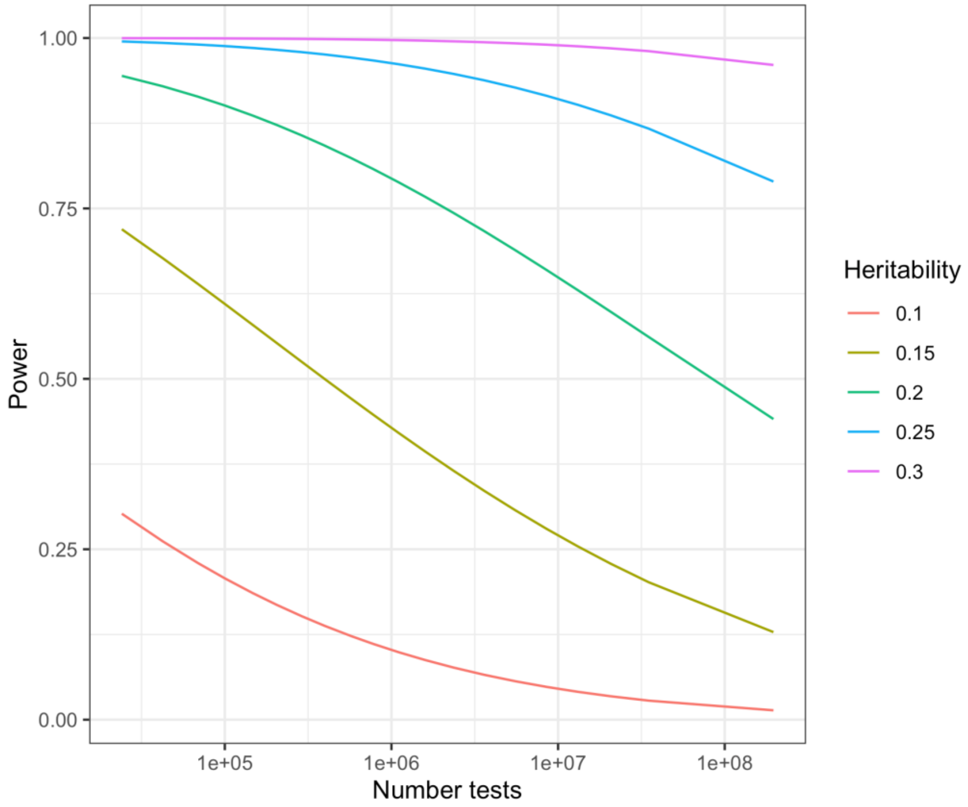


***Supplementary Figure 15.*** *Power analysis coe-QTLs. The power to detect a co-eQTL with a certain heritability is calculated based on a Bonferroni-corrected significant threshold of 0.05 and a sample size of 173. The multiple testing correction is strongly affected by the number of tests on the x-axis. The maximum number of tests in the plot represents testing all genes against each other that are expressed in monocytes for the Oelen v3 data set, but testing only one SNP per pair.*

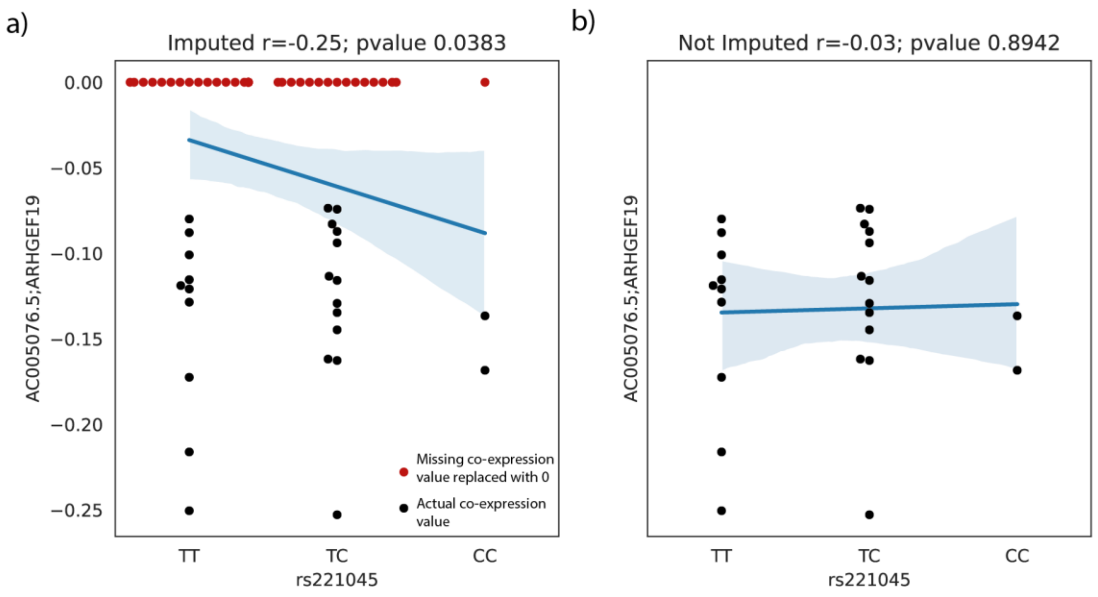


***Supplementary Figure 16.*** *Effect of replacing NaN values for the coeQTL analysis.* ***a)*** *The false positive co-eQTL identified if we replaced the NaN values to zeros.* ***b)*** *The absence of co-eQTL effects if we did not replace the NaN values to zeros.*

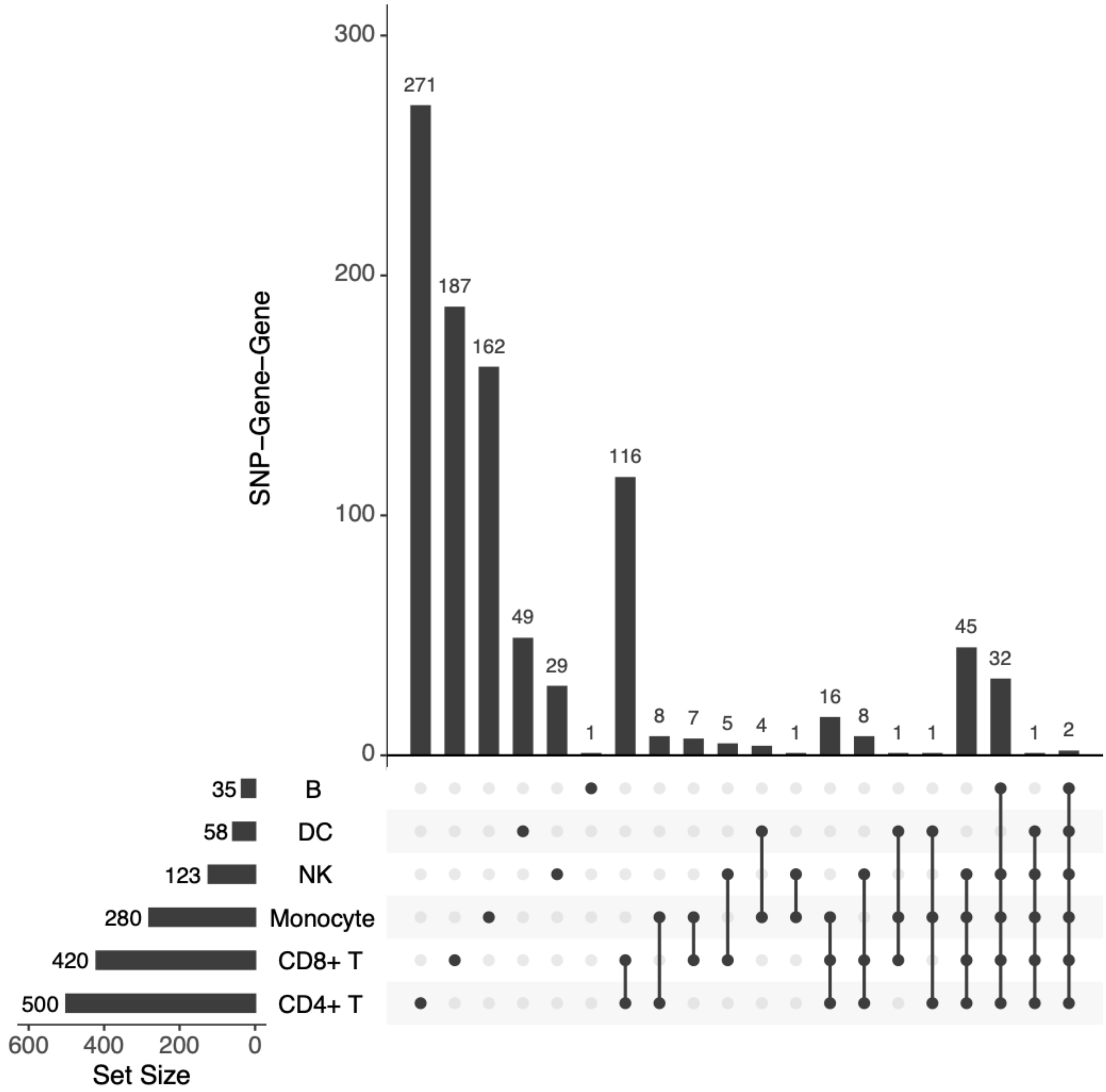


***Supplementary Figure 17.*** *Upset plot with overlap of significant co-eQTLs between cell types.*

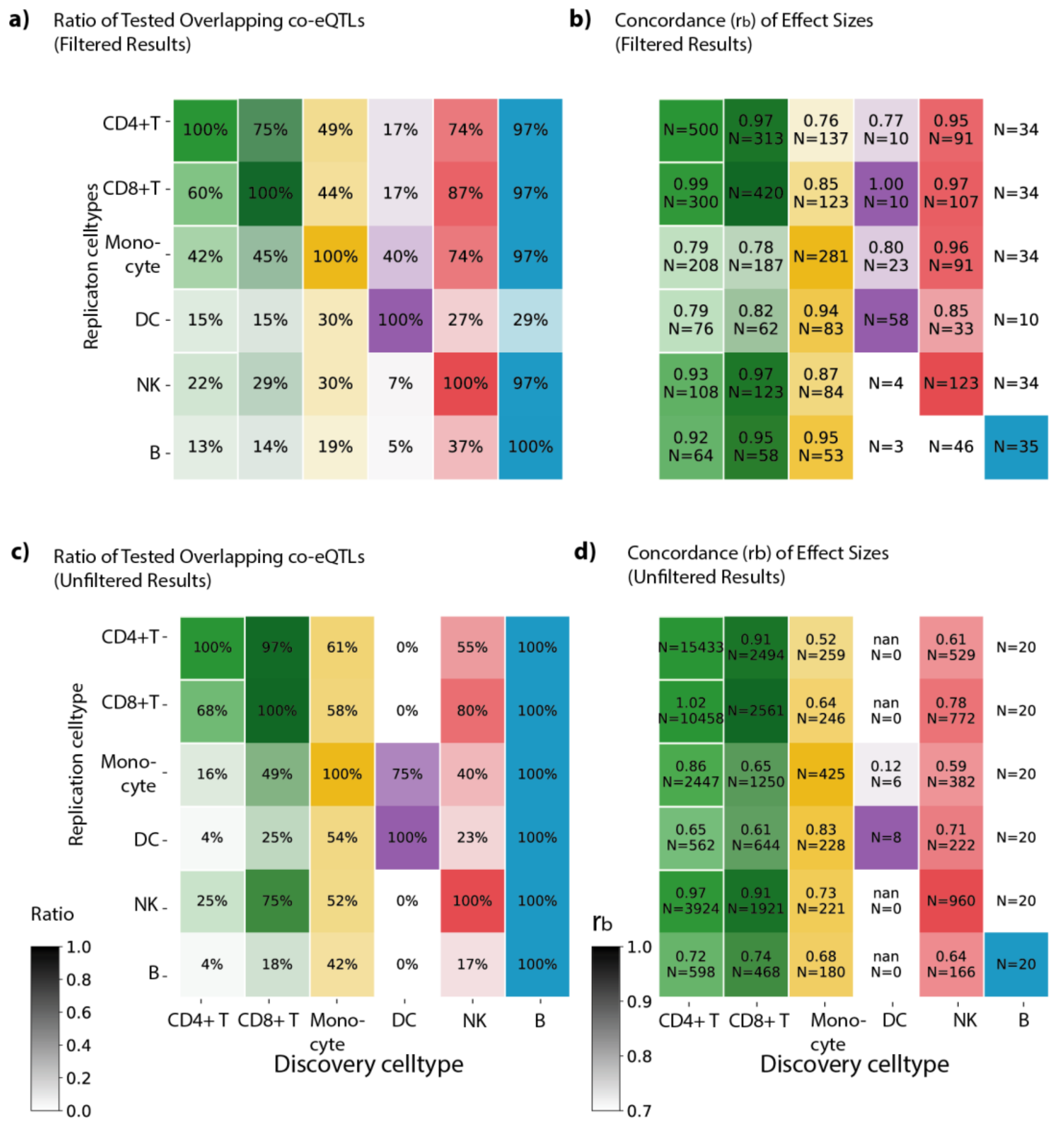


***Supplementary Figure 18****. The cell type specificity of co-eQTLs identified in each cell type. Here we replicated the identified co-eQTLs from each cell type in other cell types and used two measures to show the replication performance: the ratio of tested co-eQTLs in replication, and the concordance of effect sizes shown with the rb values. Panel* ***a)*** *shows the ratio of tested co-eQTLs in replications for the identified co-eQTLs identified with the filtering strategy. Panel* ***b)*** *shows the rb values for co-eQTLs identified with the filtering strategy. Panel* ***c)*** *shows the ratio of tested co-eQTLs in replications for the identified co-eQTLs identified without the filtering strategy. Panel* ***d)*** *shows the rb values for co-eQTLs identified without the filtering strategy.*

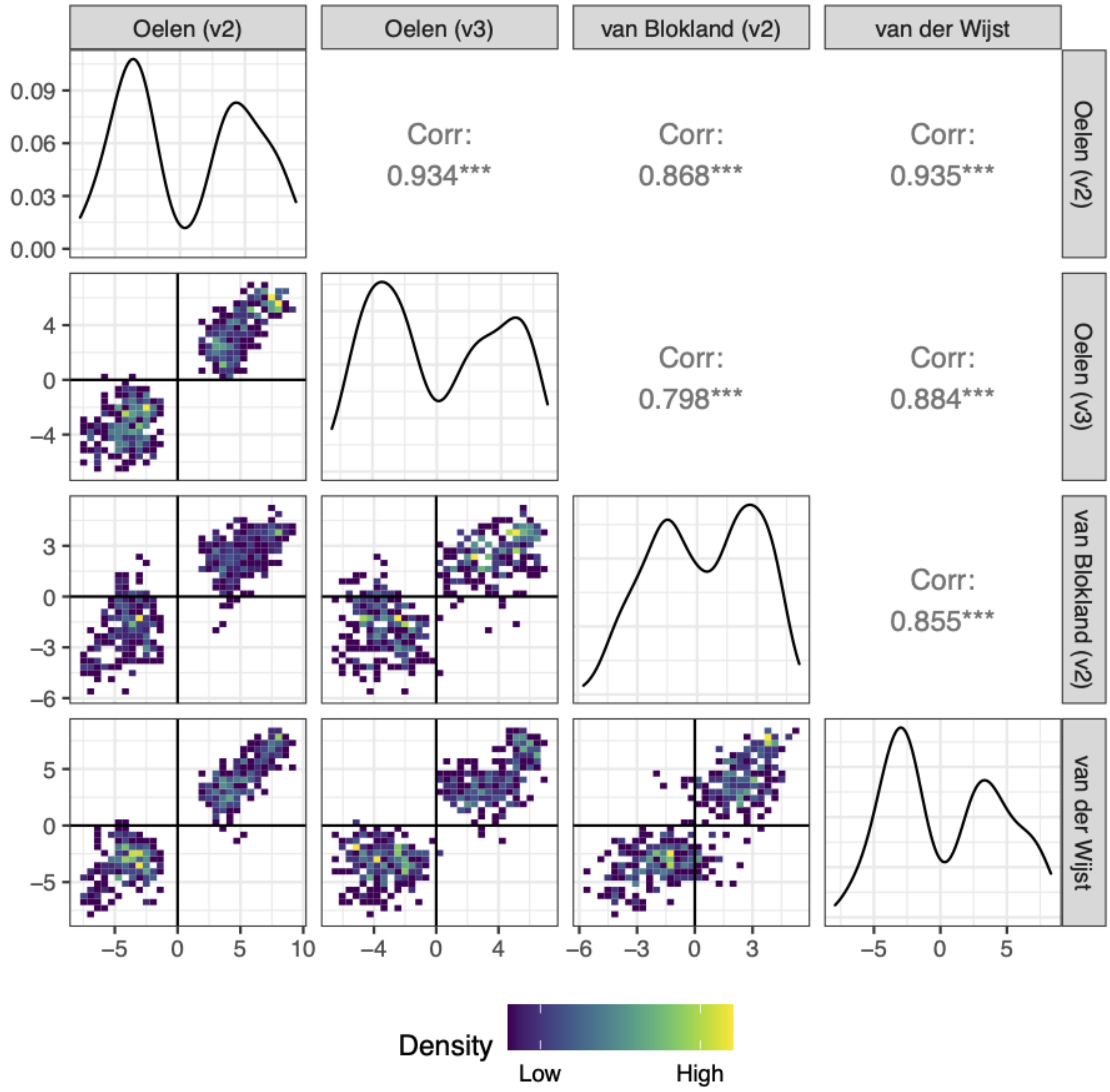


***Supplementary Figure 19.*** *Comparison of Z-scores across data sets. Distribution of significant co-eQTL Z-scores per cohort, that was included in the meta-analysis, for the CD4+ T cells. The plot shows scatter density plots between cohorts (lower triangle), distributions within the cohort (diagonal) and correlations between cohorts (upper triangle).*

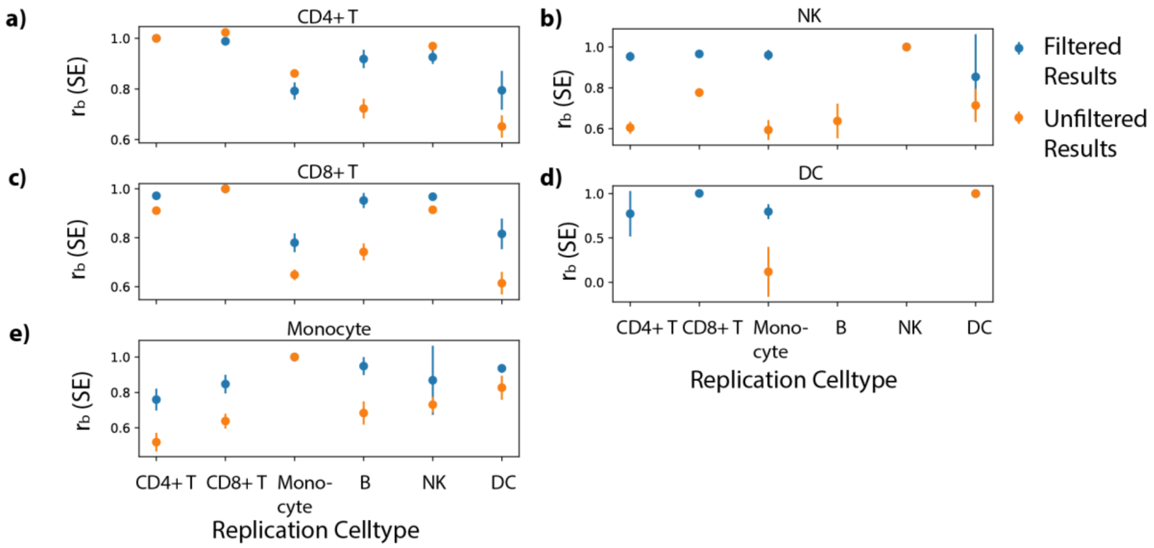


***Supplementary Figure 20**. Comparison of the rb values from the replication cell types between the co-eQTLs identified with the filtering strategy and the co-eQTLs identified without the filtering strategy.*

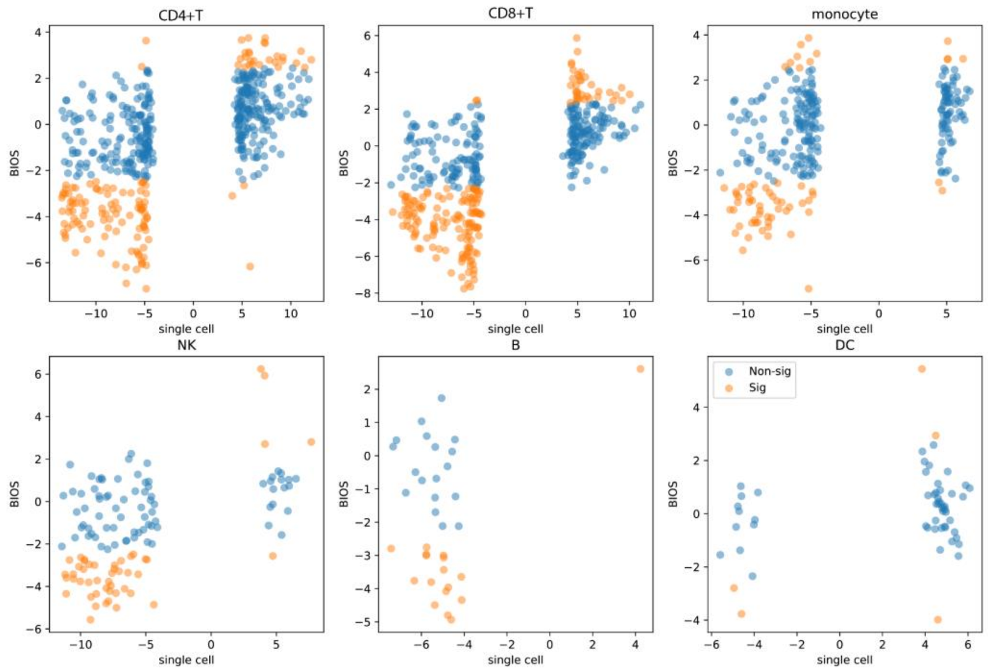


***Supplementary Figure 21.** BIOS replication for co-eQTLs identified with the filtering strategy for each cell type. The blue points labeled with "Non-sig" represent the co-*

eQTLs that could not be significantly replicated in BIOS. The orange points labeled with “Sig” represent the co-eQTLs that were replicated significantly in BIOS.

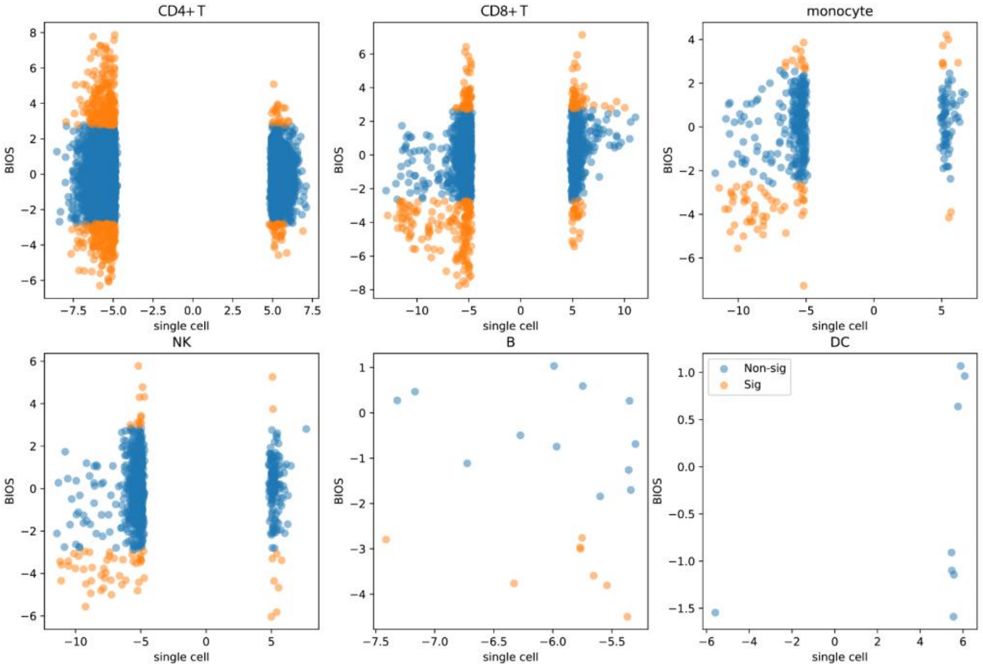


***Supplementary Figure 22.*** *BIOS replication for co-eQTLs identified without the filtering strategy for each cell type. The blue points labeled with “Non-sig” represent the co-eQTLs that could not be significantly replicated in BIOS. The orange points labeled with “Sig” represent the co-eQTLs that were replicated significantly in BIOS.*

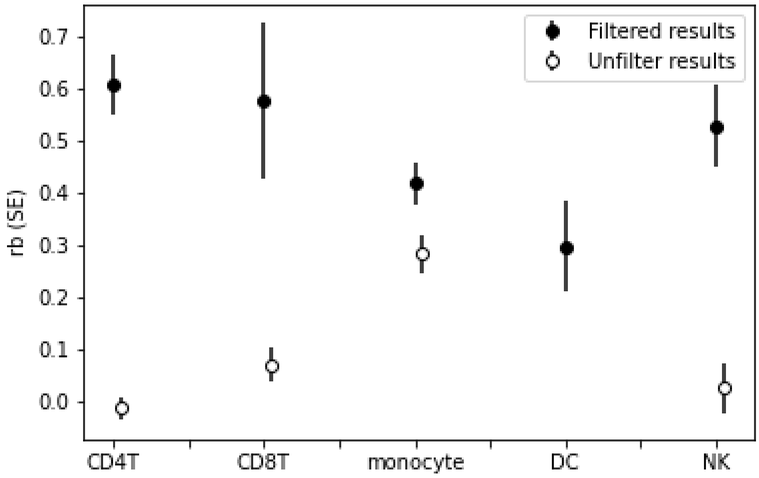


***Supplementary Figure 23.*** *Comparison of rb values from BIOS replication analysis between co-eQTLs identified with the filtering strategy and that without the filtering strategy.*

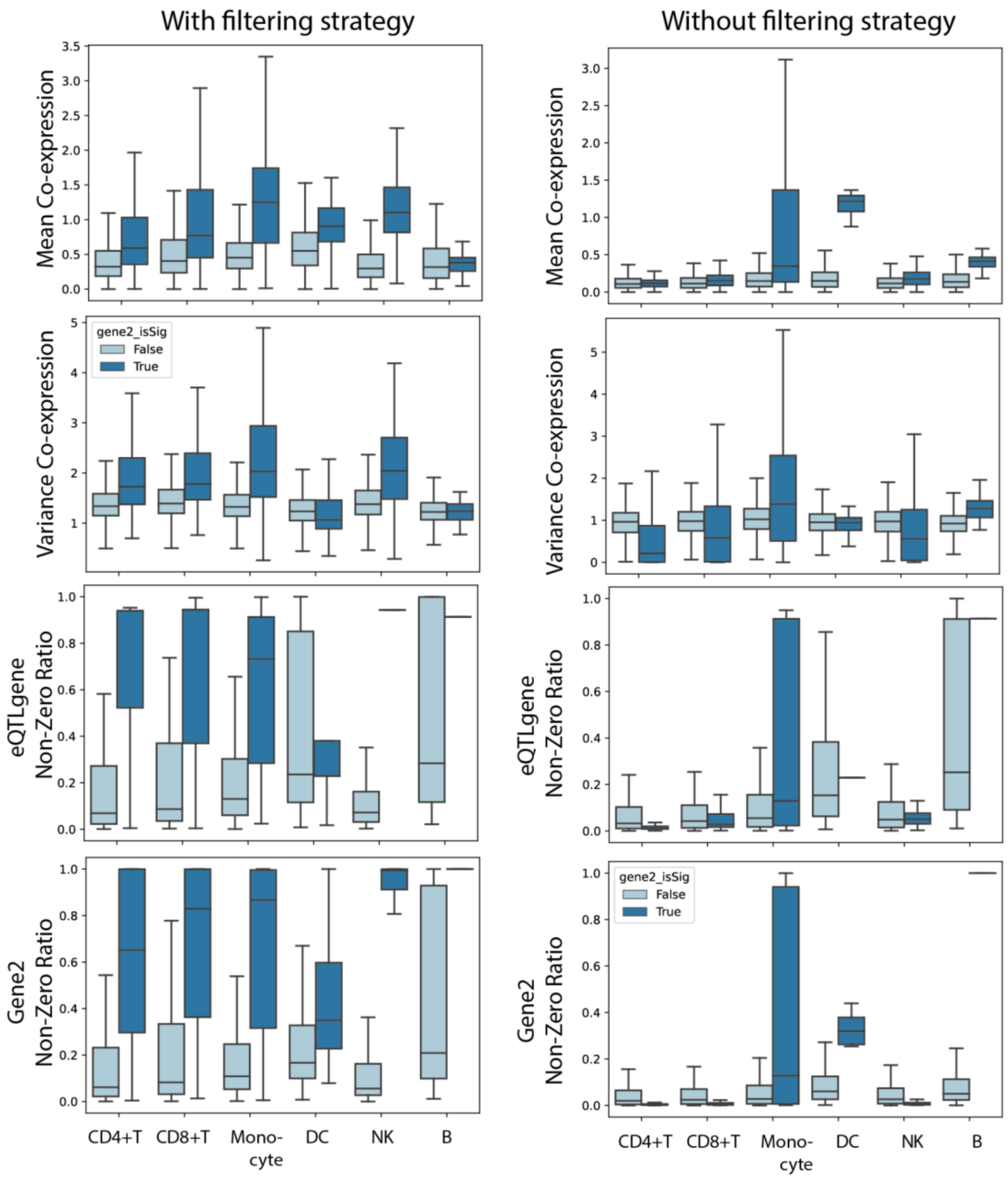


***Supplementary Figure 24.*** *Comparison between the co-eQTLs obtained with and without the filtering strategy. Panel* ***a, c, e, g*** *show values from the co-eQTLs obtained with the filtering strategy. Panel* ***b, d, f, g*** *show values from the co-eQTLs obtained without the filtering strategy. Panel a and b shows the comparison of co-expression mean values among individuals from the Oelen v2 data for different cell types between the significant co-eQTLs and the insignificant SNP-eQTLgene-gene2 triplets. Panel c and d shows the comparison of co-expression variances among individuals from the*

*Oelen v2 data for different cell types between the significant co-eQTLs and the insignificant SNP-eQTLgene-gene2 triplets. Panel e and g shows the comparison of eQTL gene non-zero ratio (the percentage of cells where this gene is expressed) from the Oelen v2 data for different cell types between the significant co-eQTLs and the insignificant SNP-eQTLgene-gene2 triplets.*

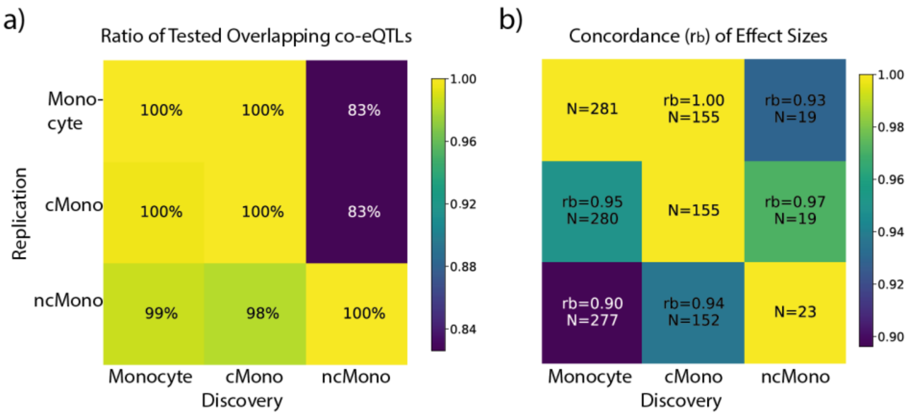


**Supplementary Figure 25.** *Impact of subcell types. Here we showed the replication performance of co-eQTLs identified with the filtering strategy in Monocytes, classical monocytes (cMono) and non-classical monocytes (ncMono) in each other. Panel* ***a)*** *shows the ratio of tested co-eQTLs in the replications. Panel* ***b)*** *shows the rb values for each replication.*

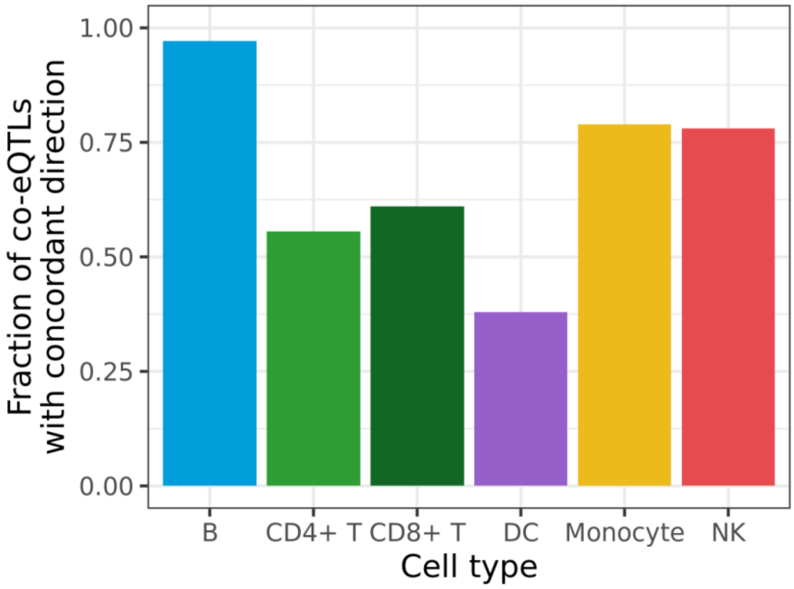

***Supplementary Figure 26.*** *Fractions of co-eQTLs in each cell type that have a concordant direction of effect compared to the associated eQTL (i.e. the correlation increases when the expression of the eGene increases).*

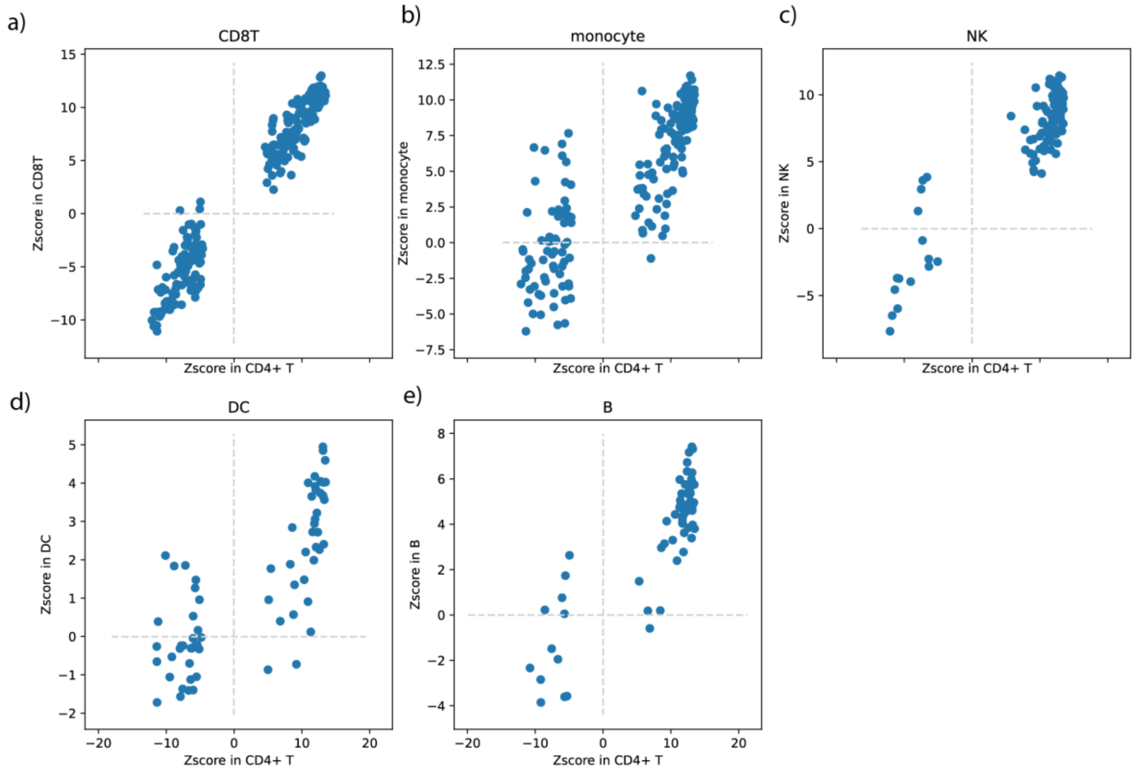


***Supplementary Figure 27.*** *The replication results for co-eQTLs identified in CD4+ T cells in other cell types. Each panel shows the replication performance in the corresponding cell type as indicated in the panel titles.*

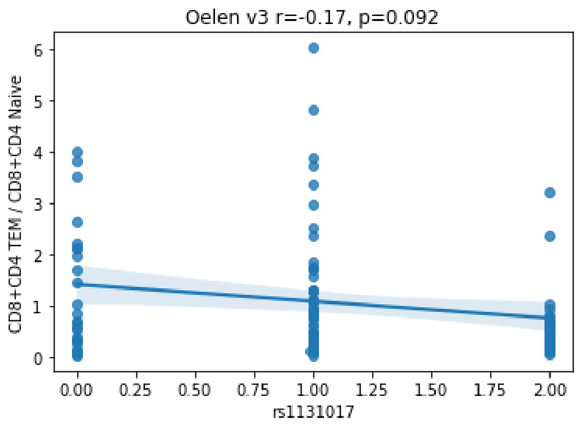

***Supplementary Figure 28****. The association between SNP rs11311017 and the ratio between CD4+ & CD8+ TEM / CD4+ & CD8+ Naive T cells*

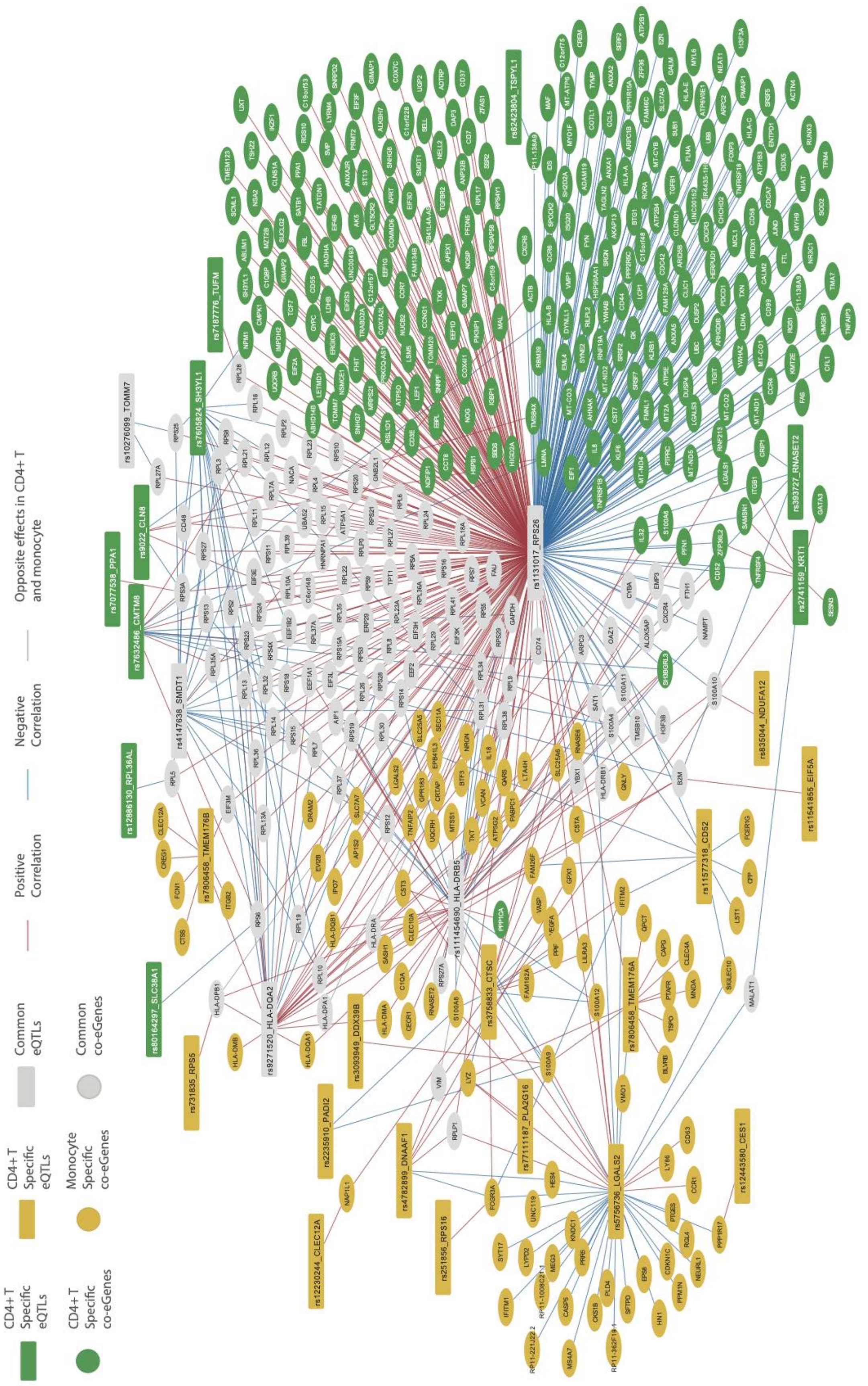


***Supplementary Figure 29**. Complete CD4+ T and Monocyte co-eQTL network*

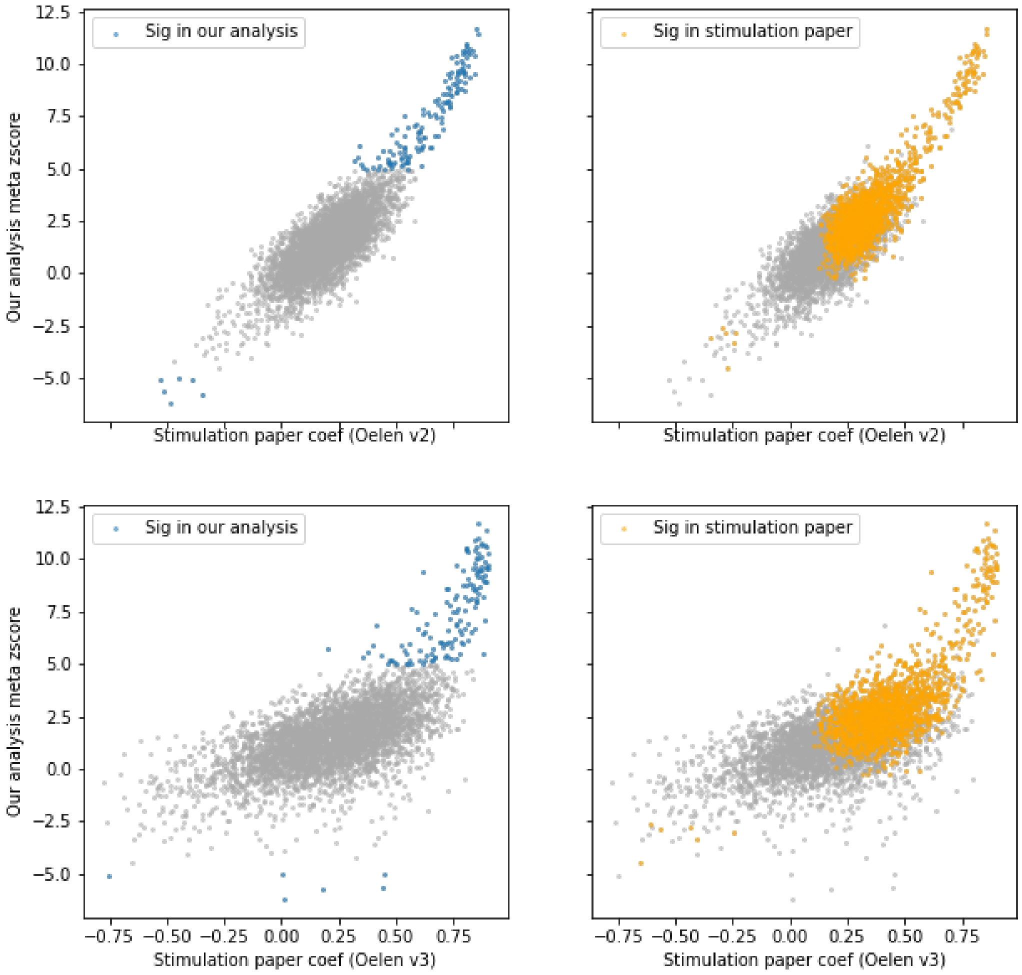


***Supplementary Figure 30***. *Correspondence between co-eQTLs from our study and co-eQTLs from Oelen et al. Here we compared the Z-scores for co-eQTLs identified for the SNP-eGene pair rs1131017 - RPS26. The blue dots indicate the significant co-eQTLs identified in our study, the orange dots indicate the significant co-eQTLs identified in the Oelen study. In total, Oelen study identified 1,564 co-eGenes, while we identified 148 in monocytes, and 91% of them were also identified with concordance effect direction as the top 10% significant outcomes in the Oelen study.*

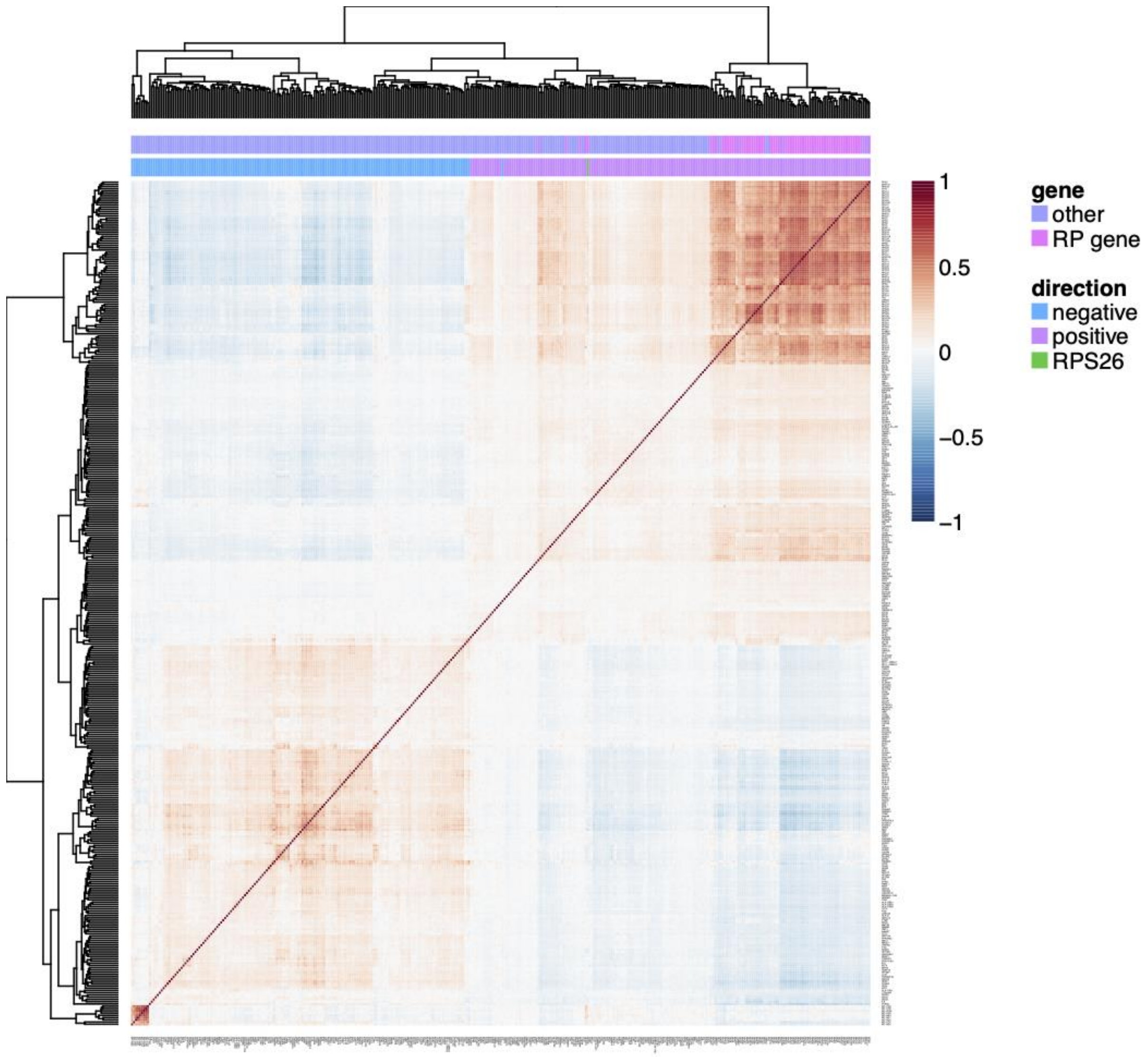


***Supplementary Figure 31****. Correlation structure of rs11311017-RPS26 co-eGenes significant in CD4+T cells. The bars above the correlation heatmap show if the gene is a ribosomal gene (RP gene) and which direction of effect the corresponding co-eQTL with this co-eGene has.*

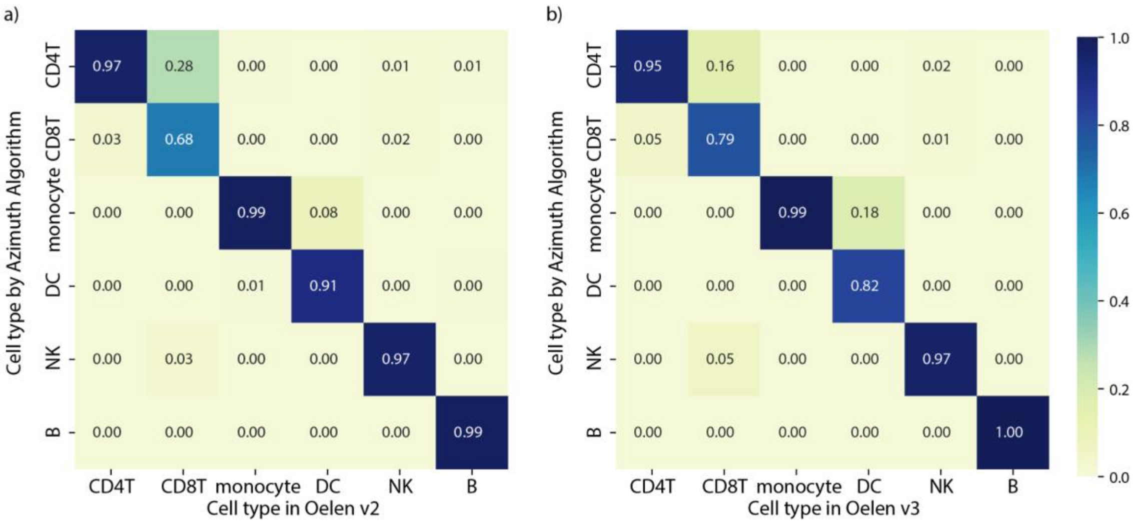


***Supplementary Figure 32.*** *Comparison between the Azimuth cell type classification and the cell type classification provided in the original publications in* ***a)*** *Oelen v2 dataset* ***3)*** *Oelen v3 dataset.*
